# Targeting TPO/MPL Signaling to Mitigate JAK2V617F-driven Cardiac Microvascular Disease

**DOI:** 10.64898/2026.04.01.715884

**Authors:** Xiaoxi Yang, Kyla Masarik, Xiaochuan Sun, Fangfang Zhang, Kirsten Zheng, Haoyi Zheng, Huichun Zhan

**Author notes:** Correspondence: Haoyi Zheng, M.D., Cardiac Imaging, The Heart Center, Saint Francis Hospital, Roslyn, NY, Correspondence: Huichun Zhan, M.D., Division of Hematology-Oncology, Department of Medicine, Stony Brook University, Stony Brook, NY 11794; Northport VA Medical Center, 79 Middleville Road, Northport, NY. **Data sharing statement:** For original data, please contact.

## Abstract

**Background:** Individuals with *JAK2V617F*-mutant myeloproliferative neoplasms or clonal hematopoiesis of indeterminate potential have a markedly increased risk of cardiovascular disease, yet the mechanisms by which mutant blood cells drive vascular and cardiac dysfunction remain incompletely understood. Although the thrombopoietin (TPO) receptor MPL is central to hematopoiesis and is expressed in vascular endothelial cells (ECs), its role in JAK2V617F-associated cardiovascular complications is unknown.

**Methods and Results:** We generated chimeric mice with *JAK2V617F*-mutant blood cells and wild-type endothelium by bone marrow transplantation and challenged them with a high-fat/high-cholesterol diet to model cardiometabolic stress. These mice developed a distinct cardiovascular phenotype characterized by microvascular disease, increased left ventricular mass, and relatively preserved left ventricular ejection fraction. Histological analysis revealed coronary arteriole stenosis, perivascular fibrosis, reduced microvascular density, and endocardial injury, without evidence of epicardial coronary stenosis or myocardium infarction.

Single-cell RNA sequencing revealed activation of inflammatory, stress-response, and endothelial-to-mesenchymal transition gene signatures in ECs, most prominently within the endocardial ECs. Immunohistochemistry identified MPL expression predominantly in endocardial ECs. TPO/MPL signaling was upregulated in endocardial ECs in mice with *JAK2V617F*-mutant hematopoiesis, and treatment with an anti-MPL neutralizing antibody markedly improved cardiac pathology, restored endocardial integrity, and increased coronary microvascular density despite persistent systemic inflammation.

**Conclusions:** *JAK2V617F*-mutant hematopoiesis induces coronary microvascular dysfunction. Endocardial ECs represent a key cellular target under cardiometabolic stress, and endocardial MPL signaling constitutes a potential targetable pathway in JAK2V617F-associated cardiovascular disease.

**Graphic Abstract:** 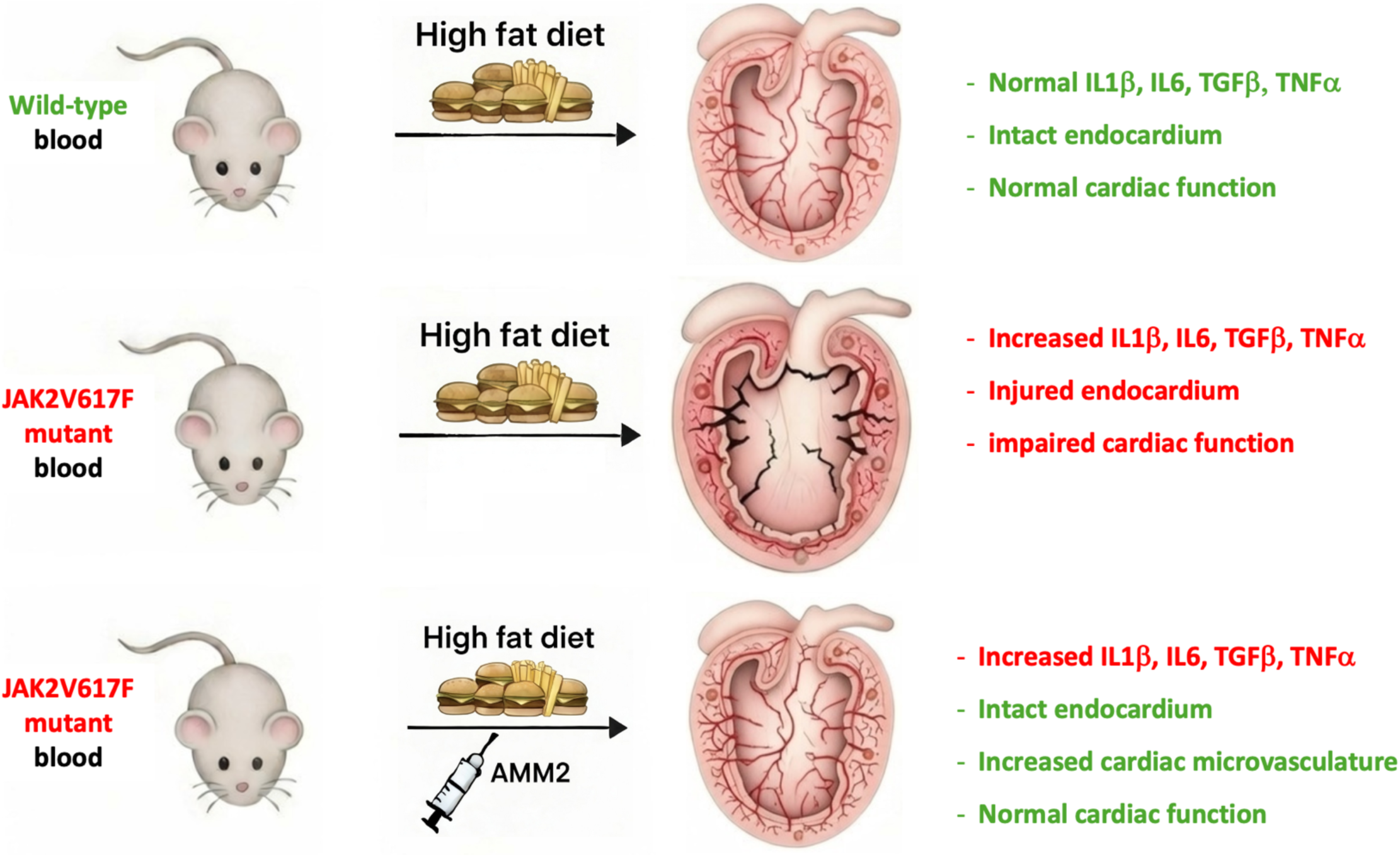

**Key Points:**

- JAK2V617F-mutant hematopoiesis induces cardiac microvascular disease
- MPL is expressed in endocardial ECs and MPL inhibition restores endocardial integrity and improves cardiac microvascular function

## Introduction

The acquired kinase mutation JAK2V617F drives clonal expansion of hematopoietic stem/progenitor cells and overproduction of mature, often dysfunctional blood cells. While it is best known as the driver mutation in myeloproliferative neoplasms (MPNs), accumulating evidence shows that JAK2V617F itself — rather than overt MPN — is a potent risk factor for cardiovascular disease (CVD). In patients with JAK2V617F-positive MPNs, CVD is the leading cause of morbidity and mortality, with arterial or venous thrombosis affecting 40-50% of patients during the disease course, far exceeding age-matched controls^1^. JAK2V617F also occurs in clonal hematopoiesis of indeterminate potential (CHIP) without overt hematologic disease, present in ∼0.1-0.2% of middle-aged populations^2^. Among all CHIP variants, JAK2V617F clones expand most rapidly^3^ and confer ∼12-fold increased risk of coronary heart disease and ischemic stroke compared to those without any CHIP mutations —substantially higher than other CHIP mutations^4,5^. Even small JAK2V617F clones (with allele burdens <2%) are associated with >5-fold increase in cardiovascular events^6^, underscoring the profound vascular impact of this mutation. Recent studies further demonstrate that JAK2V617F-mutant CHIP confers a 4-fold increased risk of heart failure independent of coronary artery disease, aligning with previous clinical observations that patients with JAK2V617F-positive MPNs have more than a 2-fold increased risk of developing heart failure^7,8^. Together, these findings highlight a key unresolved question: how do JAK2V617F-mutant blood cells, whether in overt MPN or subclinical CHIP, promote vascular injury and cardiac dysfunction? Addressing this is essential for identifying modifiable pathways linking mutant hematopoiesis to CVD and for developing targeted risk-reduction strategies in this growing patient population.

Vascular endothelial cells (ECs) are central regulators of cardiovascular function. Thrombopoietin (TPO) and its receptor MPL are best known for their roles in hematopoietic stem cell survival and megakaryocyte differentiation^9^. Although MPL is mainly expressed on hematopoietic stem cells, megakaryocytes, and platelets^10^, it is also present on vascular ECs —including human umbilical cord ECs^11,12^, and mouse liver^13^ and pulmonary^14^ ECs — where TPO/MPL signaling modulates vascular function^12,13,15–17^. While JAK2V617F is primarily expressed in hematopoietic cells in MPNs, it is also detectable in vascular ECs in some patients^18–20^. We previously showed that mice expressing JAK2V617F in both hematopoietic cells and ECs develop spontaneous heart failure with thrombosis, vasculopathy, and reduced left ventricular ejection fraction (LVEF)^21^. Using EC-specific JAK2V617F mice and MPN patient-derived ECs, we demonstrated that JAK2V617F induces endothelial-to-mesenchymal transition (EndMT)^22^ — a process normally involved in heart development but can be reactivated in postnatal life, either as an adaptative response to new physiological demands or as a maladaptive response in disease^23–26^. Notably, MPL expression is upregulated in JAK2V617F-mutant ECs compared to wild-type ECs^22,27,28^, and genetic ablation of endothelial MPL suppresses JAK2V617F-induced EndMT and prevents CVD development^22^. Because JAK2V617F-mutant hematopoiesis is far more prevalent than endothelial mutation, we asked whether mutant blood cells alone can drive cardiovascular dysfunction and whether pharmacological blockade of TPO/MPL signaling with the neutralizing antibody AMM2^29–34^ can prevent these effects. In this study, we show that JAK2V617F-mutant blood cells induce endothelial dysfunction, evidenced by transcriptomic and histologic abnormalities, even in the absence of detectable cardiac impairment. When challenged with a high-fat/high-cholesterol diet (HFD), however, these mice developed a distinct CVD phenotype characterized by cardiac microvascular disease with coronary arteriole stenosis, perivascular fibrosis, and endocardial disruption, accompanied by striking transcriptomic alterations in endocardial ECs revealed by single-cell RNA sequencing. LV mass was increased and systolic function measured by LVEF was relatively preserved. We further discover that MPL expression is selectively expressed in endocardial ECs, whereas arterial, venous, and capillary ECs were largely negative. Importantly, treatment with the MPL-neutralizing antibody AMM2 markedly ameliorates these diet-induced cardiovascular pathology, restores endocardial integrity, increases coronary vascular density, and suppresses inflammatory and stress-response gene signatures in endocardial ECs despite persistent systemic inflammation. Together, these findings suggest that endocardial MPL signaling may represent a therapeutic target to mitigate cardiovascular risk associated with JAK2V617F-mutant hematopoiesis.

## Methods

### Experimental mice

JAK2V617F Flip-Flop (FF1) mice (which carry a Cre-inducible human JAK2V617F gene driven by the human JAK2 promoter; JAX stock #037558), Tie2-Cre mice (JAX stock #008863), and CD45.1 congenic mice (JAX stock #002014) were obtained from the Jackson Laboratory. FF1 mice were crossed with Tie2-Cre mice to express JAK2V617F in hematopoietic cells, generating Tie2-cre^+^FF1^+^ mice on a CD45.2 background.

### Isolation and culture of murine cardiac endothelial cells (ECs)

Primary murine cardiac ECs (CD45^-^CD31^+^) were isolated from mouse heart following a previously described protocol with minor modification^21,22,27,35,36^. Briefly, mice were euthanized, and the thoracic cavity was immediately opened via a midline sternotomy. Cardiac perfusion was performed with 20 ml cold PBS using a 25-gauge needle inserted into the left ventricular apex, with concurrent drainage through a small incision in the right atrium. The heart was collected, rinsed three times in PBS, and finely minced with scissors. The tissue fragments were then digested in DMEM medium containing 1 mg/mL Collagenase/Dispase (Roche, Switzerland) and 25 U/mL DNase (Sigma) at 37℃ for 1 hour with gentle shaking. The digested tissue was then triturated using a 20ml syringe attached to a 16-gauge needle for at least 12 passes to obtain a single-cell suspension, filtered through a 70µm cell strainer, washed with 40ml PBS, and centrifuged at 500g for 10 minutes. The resulting cell pellet was resuspended and sequentially processed for CD45^+^ cell depletion followed by CD31^+^ positive selection using magnetically labeled microbeads (Miltenyi Biotec, San Diego, CA) according to the manufacturer’s protocol. Purified ECs (CD45^-^CD31^+^) were cultured on 2% gelatin coated plates in complete EC medium^35^. No medium change was performed for the first 72 hours to allow EC attachment, followed by medium change every 2-3 days.

### In vitro cell culture

For EC co-culture with blood mononuclear cells, primary murine cardiac ECs (2×10^4^) were seeded onto 1% gelatin-coated 24-well plates and cultured in 500ul of mouse complete EC medium starting on Day −2. On Day 0, ECs were co-cultured with 4×10^4^ JAK2 wild-type blood mononuclear cells (isolated from control mice) or JAK2V617F mutant blood mononuclear cells (isolated from Tie2-cre^+^FF1^+^ mice) placed in Transwell inserts (1.0um pore size; Corning, NY) containing complete EC medium. After two days of co-culture (Day 2), ECs were harvested, and their *in vitro* angiogenic capacities were assessed.

For AMM2 treatment experiment, primary murine cardiac ECs (2×10^4^) were seeded onto 2% gelatin-coated 24-well plates and cultured in 500ul of complete EC medium starting on Day −2. On Day 0, ECs were treated with either phosphate buffered saline (PBS) or 100ng/mL AMM2, an MPL-neutralizing antibody (Cat. #10401, IBL-America, Minneapolis, MN), for 48 hours. On Day 2, ECs were harvested, and their *in vitro* angiogenic capacities were assessed.

### Endothelial cell in vitro angiogenesis assay

The EC tube formation assay was performed to evaluate in vitro angiogenic capacity^21,22,27,28,37^. Matrigel® matrix (Cat. 354234, Corning) was thawed overnight at 4°C and kept on ice until use. A pre-chilled 48-well culture plate was coated with 150μl of Matrigel per well and incubated at 37°C for 30 minutes to allow gelation. 6 × 10^4^ primary mouse heart ECs in 300ul of EC medium were seeded onto the solidified Matrigel. Tube formation was imaged at 2, 4, 6, 8, and 24 hours using a phase-contrast microscope (AMEX-1200, ThermoFisher) at 4x and 10x magnifications. Quantitative analysis was performed using the ImageJ® Angiogenesis Analyzer^38^ (National Institute of Health, Bethesda, MD) by counting the number of tubes and nodes in across twenty to thirty non-overlapping 400 x 400 pixel fields.

### Stem cell transplantation, diet, and AMM2 treatment

Approximately 1×10^6^ marrow cells (either JAK2 wild-type cells from control mice or JAK2V617F mutant cells from age-matched Tie2-cre^+^FF1^+^ mice; CD45.2 background) were transplanted into lethally irradiated (two 540cGy doses, 3 hours apart) 8-week-old wild-type recipient mice (CD45.1) via intravenous tail vein injection. Peripheral blood was collected every 4 weeks after transplantation to monitor complete blood counts and CD45.2^+^ donor chimerism.

For diet and AMM2 treatment: eight weeks after transplantation, mice were assigned to one of the following groups for five weeks: (1) Standard diet: PicoLab® Rodent Diet 20 5053; (2) High-fat diet (HFD): TD88137 (Inotiv, Indianapolis, IN), containing 21% (wt/wt) milk fat and 49% (wt/wt) carbohydrate (sucrose and corn starch); (3) HFD plus AMM2 treatment: AMM2 (IBL-America #10401, Minneapolis, MN) administered intravenously at 0.1mg/kg every 10 days for a total of four doses.

Mice were assigned to experimental groups using stratified randomization to ensure comparable male-to-female ratios across groups. Echocardiography was performed in a semi-blinded manner as the sonographer measured a large number of animals and was not aware of individual treatment assignments at the time of data acquisition.

### Complete blood counts

Peripheral blood was obtained from the facial vein via submandibular bleeding and collected in EDTA coated tubes. The samples were then analyzed using a Vetscan Hm5 Hematology Analyzer (Abaxis) to determine the blood count.

### Transthoracic echocardiography

Transthoracic echocardiography was performed in mildly anesthetized, spontaneously breathing mice maintained under 1-3% isoflurane in oxygen at a flow rate ∼1L/min using a Vevo 3100 high-resolution imaging system (VisualSonics Inc, Toronto, Canada), as previously described^21,22^. Parasternal long-axis and sequential parasternal short-axis views were acquired to assess global and regional wall motion. Left ventricular (LV) end-systolic and end-diastolic dimensions were measured at the mid-ventricular level, and LV ejection fraction and fractional shortening were measured using standard formulas to evaluate systolic function^39^.

### Histology

Hearts were fixed in cold 4% paraformaldehyde overnight at 4°C with gentle shaking, then washed several times with PBS at room temperature to remove residual paraformaldehyde. Paraffin-embedded sections (5μm thick) were stained with hematoxylin/eosin (H&E) or Masson’s trichrome using standard protocols and reagents from Sigma (St. Louis, MO). Whole-slide images were acquired using a NanoZoomer S20 Digital Slide Scanner (Hamamatsu Photonics K.K., Japan), and representative images were captured with a Nikon Eclipse Ts2R Inverted Microscope.

Coronary microvasculopathy was assessed on H&E-stained images using Image J (NIH) as previously described^22^. The ratio of luminal radius to wall thickness was quantified for each coronary arterioles (10-100um in diameter), averaging four different measurement (top-bottom, left-right, and both diagonals). Arterioles were considered stenotic when the luminal radius-to-wall thickness ratio was <1^40,41^.

Perivascular and subendocardial collagen content was quantified on Masson’s Trichrome-stained sections, where collagen fibers appear blue, muscle fibers red, and nuclei black. For perivascular collagen quantification, intramyocardial vessels with circular profiles and intact morphology (10–100 µm in diameter) were selected. The collagen-positive (blue) area surrounding each vessel was measured in ImageJ and expressed as a percentage of the total vessel area. For subendocardial collagen quantification, collagen content beneath the endocardial surface of both left and right ventricles was calculated as the ratio of collagen-positive subendocardial area to the length of the corresponding endocardial segment.

### Single-cell RNA sequencing and data analysis

Single-cell suspensions from isolated mouse hearts were prepared by tissue mincing and enzyme digestion as described above in “*Isolation and culture of murine cardiac endothelial cells*”, with the exception that the digestion step was shortened to two 15-minute incubations (30 minutes in total). The digested tissue was gently triturated using a 20ml syringe attached to a 16-gauge needle for at least 12 passes to obtain a single-cell suspension, filtered through a 40µm cell strainer, washed with 40ml PBS containing 2%FBS, and centrifuged at 500g for 10 minutes. Red blood cells were lysed one to two times using pre-warmed Red Blood Cell Lysing Buffer (Cat. R7757, Sigma, Germany) with a 5-minute incubation at room temperature. After washing, cell pellets were resuspended in 100ul of Dead Cell Removal Microbeads per 10^7^ total cells (Cat. 130-090-101, Miltenyi Biotec) and incubated 15 minutes at room temperature. The mixture was diluted in 1x binding buffer to a minimum volume of 500ul and applied to a Large cell column (Cat. 130-042-202, Miltenyi Biotec) mounted on a MACS Separator (Cat. 130-042-602, Miltenyi Biotec, Germany) to deplete dead cells bound to magnetic beads. Cells were washed twice with PBS containing 0.04% bovine serum albumin (BSA) and resuspended in PBS/ 0.04% BSA at a final concentration of 0.7-1.2 x 10^6^ cells/mL.

Freshly isolated single-cell suspensions were immediately processed at the Single Cell Core Facility to generate single cell RNA-sequencing libraries using the Chromium Next GEM Single Cell 5′ Reagent Kit v2 (10x Genomics, Pleasanton, CA) according to the manufacturer’s instructions. Briefly, cells, gel beads, and partitioning oil were loaded into a Chromium Next GEM Chip G for a target recovery of 10,000 cells per sample. Gel Bead-in-Emulsions (GEMs) were generated on a Chromium Controller (10x Genomics). Following reverse transcription using a GEM–RT incubation protocol, the GEMs were broken with Recovery Agent (10x Genomics), and cDNA was purified using DynaBeads MyOne Silane beads (10x Genomics), amplified by PCR (13 cycles), and purified using SPRIselect reagent (Beckman Coulter, Brea, CA, USA). cDNA concentration and amplicon size were measured using a TapeStation 4200 with a D5000 high sensitivity ScreenTape system (Agilent, Santa Clara, CA, USA). After fragmentation, end-repair, A-tailing, and size selection, cDNA was ligated with adaptors and purified again using SPRISselect reagent, according to 10x Genomics instructions. Libraries were amplified by PCR (10-11 cycles) using indexed oligonucleotide primers from the Chromium i7 Multiplex Kit (10x Genomics). The final libraries underwent double-sided size selection with SPRISselect reagents and were quality-checked on a TapeStation 4200. cDNA libraries were sequenced on an NovaSeq XPlus (Illumina) using paired-end 150 reads (Read 1: 151bp; i7 index: 10bp; Read 2: 151bp) at Novogene (Durham, NC, USA).

Raw sequencing data FASTQ files were aligned to the mm10-2020-A mouse reference genome and processed with CellRanger 7.1.0 (10x Genomics). Downstream analysis was performed using the Seurat 5.2.0^42^. Ambient RNA contamination was corrected using SoupX^43^, and doublets were identified and removed using DoubletFinder^44^. Cells were filtered to include those with ≥300 detected genes (nGenes), ≥500 unique molecular identifiers (nUMI), <15% mitochondrial gene content, and a log-normalized genes-to-UMI ratio > 0.8. Data were normalized using NormalizeData with default parameters, and the top variable genes were identified using FindVariableFeatures (selection.method = “vst” and features = 3,000). After scaling (ScaleData), dimensionality reduction was performed by principal component analysis (PCA), using top 20 PCs as determined by an Elbow Plot. Data integration and clusters were visualized using a 2D Uniform Manifold and Approximation Projection (UMAP). Cluster annotation was based on canonical marker genes identified from published literature. Marker expression was visualized in both UMAP and violin plots. Differentially expressed genes were identified using the FindMarkers function (Seurat) with adjusted p-value ranking. Gene set enrichment analysis (GSEA) was performed using GSEA v4.4.0^45^.

### Reverse transcription polymerase chain reaction (RT-PCR) assays

CD45^-^CD31^+^ ECs were isolated from marrow, spleen, lung, and heart tissue single cells using magnetically labeled microbeads (Miltenyi Biotec, San Diego, CA) according to the manufacturer’s protocol^21,22,27,35,36^. Total RNA was isolated using the RNeasy mini kit (Qiagen) and was digested with RNase-free DNase (Qiagen). The RNA was then reverse transcribed to cDNA using the high-capacity cDNA reverse transcription kit (Thermo Fisher) following the manufacturer’s protocol. The TaqMan® Gene Expression Assay (Mm00440306_g1, ThermoFisher) was used to detect mouse MPL gene expression in ECs.

### Flow cytometry

Cells were counted and stained with antibodies by incubating cell suspensions at 4°C for 30 minutes in the dark. Samples were analyzed using a CytoFLEX flow cytometer (Beckman Coulter, Indianapolis, IN, USA). CD45 (Clone 104, Biolegend, San Diego, CA, USA), CD45.2 (Clone 104, Biolegend), CD45.1 (Clone A20, BD Biosciences, San Jose, CA), Gr-1 (Clone RB6-8C5, Biolegend), CD11b (Clone M1/70, Biolegend), CD45R/B220 (Clone RA3-6B2, Biolegend), CD3 (Clone 145-2C11, Biolegend), CD41 (Clone MWReg30, Biolegend), Lineage (Lin) cocktail (include CD3, B220, Gr1, CD11b,Ter119; Biolegend), cKit (Clone 2B8, Biolegend), Sca1 (Clone D7, Biolegend), CD150 (Clone mShad150, eBioscience, San Diego, CA, USA), CD48 (Clone HM48-1, Biolegend), CD31 (Clone 390, BD Biosciences), MPL (Clone AMM2, IBL-America), and Streptavidin PE (Cat. 405203, Biolegend) antibodies were used. Flow cytometry data were analyzed using Kaluza (Beckman Coulter).

### Immunohistochemistry

Formalin-fixed, paraffin-embedded (FFPE) heart sections were baked at 60℃ for 20-30min in hybridization oven to melt the paraffin, followed by sequential washes in xylene (3, 5, and 5 minutes) and graded ethanol (100%, 95%, 70%, and 50%; 3-5 minutes each) to remove paraffin and rehydrate the tissue. Slides were then rinsed under running tap water for 5minutes.

Heat-induced antigen retrieval was performed by immersing slides in pre-heated Tris-EDTA buffer (pH 9) and heating in a steamer at 98°C for 25 minutes. Slides were cooled in the same buffer to room temperature (≥30 minutes), rinsed in TBST buffer for 5 minutes, and blocked with 1% bovine serum albumin (BSA) in TBST for 30 minutes at room temperature inside a humidity chamber.

Primary antibodies diluted in 1% BSA were applied and incubated overnight at 4°C in a humidity chamber: rabbit anti-mouse MPL (Ab232755, Abcam; 1:500), rabbit anti-mouse CD41 (Ab134131, Abcam; 1:800), rabbit anti-mouse CD31 (Ab182981; 1:2000), and mouse anti-human/mouse alpha-smooth muscle actin (aSMA, ThermoFisher; 1:500).

After removing the primary antibody, slides were washed three times in TBST (5 minutes each). Endogenous peroxidase activity was quenched with 0.6% hydrogen peroxide for 5 minutes at room temperature, followed by three additional TBST washes. Secondary antibody incubation was performed for 1 hour at room temperature using a goat anti-rabbit IgG HRP-linked antibody (Cat. 7074, Cell Signaling Technology; 1:100) diluted in 1% BSA.

Following three TBST washes, signal detection was performed using the AEC substrate kit (Abcam, ab64252) for 5 minutes for MPL, CD41, and CD31 staining and 1 minute for aSMA staining. After removing the AEC substrate, slides were rinsed twice with deionized water (3 minutes each), counterstained with Mayer’s hematoxylin (Cat. 786-1264, G-Biosciences) for 15-30 seconds, washed gently in running deionized water for 5 minutes, and treated in a bluing solution (10g magnesium sulfate (MgSO₄) and 0.67g sodium bicarbonate (NaHCO₃) in 1 L of deionized water) for 30 seconds. After a final rinse in deionized water (5 minutes), slides were mounted with aqueous mounting medium (Cat. 27290, Cell Signaling Technology) and scanned using a NanoZoomer S20 Digital Slide Scanner (Hamamatsu Photonics K.K., Japan).

### VE-cadherin in vivo staining and immunofluorescence imaging of heart vasculature

Twenty micrograms of Alexa Fluor 647-conjugated monoclonal antibody that targets mouse VE-cadherin (clone BV13, Biolegend) was injected retro-orbitally into mice under anesthesia. Ten minutes after antibody injection, the mice were euthanized. Freshly harvested hearts were fixed in 4% paraformaldehyde in PBS for 6 hours at 4°C while rotating. The tissues were washed in PBS overnight to remove the fixative and cryoprotected in a 20% sucrose PBS solution at 4 °C. The tissues were embedded in OCT (Tissue-Tek) and flash-frozen at −80°C.

To quantify coronary vascular density (VE-cadherin-positive endothelial area as a percentage of total tissue area), 5μm-thick heart sections were imaged using a Nikon Eclipse Ts2R Fluorescent Microscope with a 20x objective. Image analysis was performed using the ImageJ software (National Institute of Health). *<u>First</u>*, images were converted to 32-bit grayscale. Detection threshold was adjusted to include all tissue while excluding empty areas, and the total tissue area was measured. *<u>Next</u>*, the images were processed using the “FindEdges” function and detection threshold was adjusted consistently across images to best isolate VE-cadherin signal from background autofluorescence, and the VE-cadherin-positive area was measured. *<u>Last</u>*, vascular density was calculated as (VE-cadherin-positive area) / (total tissue area) x 100.

### Measurement of peripheral blood plasma cytokines

Peripheral blood was collected in an EDTA tubes, mixed gently by inverting the tubes 3-5 times, and immediately centrifuged at 2000g for 10 minutes at 4°C. The resulting platelet-poor plasma was collected and aliquoted and stored at −80℃ until further analysis. Cytokines levels were measured using ELISA kits following the manufacturer’s instructions: IL-1 beta (R&D Systems, Cat. MLB00C), IL-6 (R&D Systems, Cat. M6000B), TNF-alpha (R&D Systems, Cat. MTA00B), TGF-beta 1 (R&D Systems, Cat. DB100C), TPO (R&D Systems, Cat. MTP00), Platelet factor 4 (R&D Systems, Cat. MCX400). All samples were analyzed in duplicate.

### Quantification and statistical analysis

Statistical analyses were performed using GraphPad Prism 10 (San Diego, CA) and *R* software. All measurements were taken from distinct samples. Except for the scRNAseq experiments, all assays were performed independently at least twice. ANOVA was used to assess differences in means among multiple groups, while an unpaired two-tailed Student’s *t*-test was applied to compare differences between two groups. For *in vivo* experiments, randomization and blinding were not used. Sample size for animal studies were chosen to ensure a balance between the minimum numbers necessary for efficient analysis while accounting for biological variability. Figures were generated using GraphPad Prism 10, R software, and BioRender.com.

## Results

### JAK2V617F Mutant Hematopoiesis Induce Early-stage CVD in Mice on a Regular Chow Diet

To study the impact of JAK2V617F mutant blood cells on endothelial function, primary murine heart ECs (CD45^-^ CD31^+^) were isolated from wild-type mice and co-cultured with JAK2V617F mutant blood cells for 48 hours. Tube formation in Matrigel, a measure of *in vitro* angiogenesis, was mildly but significantly enhanced in wild-type ECs after co-culture with JAK2V617F mutant blood cells compared to co-culture with wild-type blood cells (Figure 1A).

**Figure 1.**
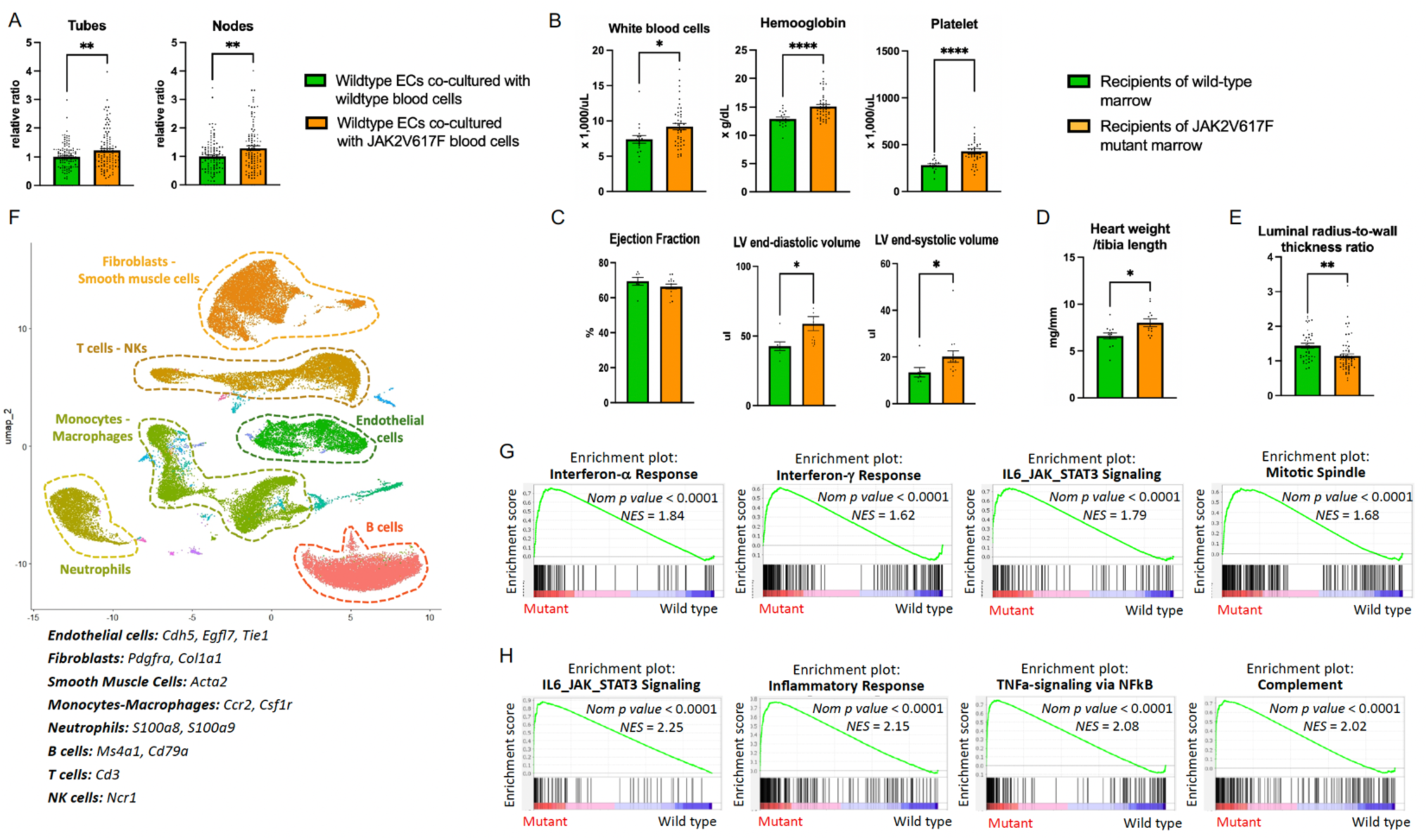
JAK2V617F mutant hematopoiesis induce early-stage CVD in mice on a regular chow diet. (**A**) Quantitative analysis showing a modest increase in tube formation in wild-type murine heart ECs co-cultured with JAK2V617F mutant blood cells compared with those co-cultured with wild-type blood cells. Data shown are representative of three independent experiments that gave similar results. (**B**) Peripheral blood counts demonstrating elevated white blood cell count, hemoglobin, and platelets in recipient mice transplanted with JAK2V617F mutant marrow compared with wild-type marrow 8 weeks post-transplant. (**C**) Transthoracic echocardiography showing preserved LV ejection fraction but increased LV end-diastolic volume and end-systolic volumes in recipients of JAK2V617F marrow compared to wildtype marrow six months after transplantation. n=7-13 mice per group from 2-3 independent experiments. (**D**) Heart weight normalized to tibia length in recipients of JAK2V617F marrow (n=12) and wild-type marrow (n=13) at 4-6 months post-transplantation. (**E**) Luminal radius-to-wall thickness ratio measured in nonoverlapping coronary arterioles from recipients of JAK2V617F marrow and wildtype marrow (7-9 mice per group; 48-93 microvessels analyzed per group). (**F**) Uniform manifold approximation and projection (UMAP) plot of unfractionated cardiac cells, with key marker genes used to define major cell types. (**G-H**) Gene set enrichment analysis (GSEA) of differentially expressed genes in cardiac ECs (**G**) and fibroblasts-smooth muscle cells (**H**) from mice with JAK2V617F mutant versus wild-type blood cells. \**P*<0.05, ** *P*<0.01, \*\*\*\**P*<0.0001. Data are presented as mean ± SEM and *P* value was calculated using an unpaired two-tailed Student’s t-test.

To assess these effects *in vivo*, we transplanted JAK2V617F-mutant bone marrow (from Tie2-cre^+^FF1^+^ mice, in which the mutation is expressed in all hematopoietic cells) or control marrow into lethally irradiated wild-type recipients, generating chimeric mice with mutant hematopoiesis and wild-type ECs. As expected, recipients of mutant marrow developed an MPN phenotype with elevated blood counts (Figure 1B). Although left ventricular (LV) ejection function remained normal over six months by serial transthoracic echocardiography (66% vs. 69%, *P*=0.261), LV end-diastolic (59ul vs. 43ul, *P*=0.013) and end-systolic (20ul vs. 13ul, *P*=0.046) volumes were increased in mutant mice compared to wild-type controls (Figure 1C). Consistently, mutant mice exhibited an increased heart weight-to-tibia length ratio at 4-6 months post-transplantation (Figure 1D). Coronary vascular analysis revealed mild arteriole narrowing in mutant mice (luminal radius-to-wall thickness ratio 1.1 vs. 1.4, *P*=0.002) (Figure 1E). In contrast to our prior murine models with combined endothelial and hematopoietic mutations, which showed prominent thrombosis in the lungs, right ventricles, and coronary arterioles^21^, no overt thrombosis was observed in mice with mutant blood cells alone.

To further investigate how mutant blood cells alter cardiac ECs, we performed single-cell RNA sequencing (scRNAseq) on cardiac cells (excluding cardiomyocytes)^46^ from recipients of mutant or wild-type blood cells ∼16 weeks post-transplantation (Figure 1F). Cardiac ECs (Cdh5^+^, Egfl7^+^, Tie1^+^) from mutant recipients showed significant enrichment of interferon-α/γ response, IL6-JAK-STAT3 signaling, and mitotic spindle pathways, consistent with endothelial inflammation and activation (Figure 1G). Taken together, these findings indicate that JAK2V617F-mutant blood cells induce early microvascular remodeling and vascular inflammation.

Before the onset of overt LV dysfunction, the heart often mounts a diffuse stress response. ECs, which are in direct contact with circulating mutant blood cells, often release cytokines and growth factors that drive fibroblast and smooth muscle cell remodeling^47^. Consistent with this, fibroblasts (Pdgfra^+^, Col3a1^+^, Col1a2^+^, Col1a1^+^) and smooth muscle cells (Acta2^+^, Tagln^+^)^46^ exhibited similar GSEA enrichment (i.e., up-regulation of IL6-JAK-STAT3 signaling, inflammatory response, TNFa-signaling via NFkB) (Figure 1H), indicating a coordinated endothelial-mesenchymal program in mice with JAK2V617F mutant blood cells. These changes were absent in monocytes/macrophages, neutrophils, T/NK cells, or B cells, arguing against artifactual enrichment due to blood cell contamination.

### JAK2V617F Mutant Hematopoiesis Promotes a Distinct Cardiac Microvascular Disease Phenotype under a High-Fat/High-cholesterol Diet

To test whether JAK2V617F mutant blood cells promote CVD under metabolic stress, we treated the chimeric mice with mutant blood cells and wild-type ECs with either a standard diet or a high-fat/high-cholesterol diet (HFD) (TD.88137; Harlan Teklad). After 5 weeks, mutant mice exhibited reduced LV ejection fraction (59% vs. 68%, P=0.001) and increased LV end-diastolic (52ul vs. 43ul, *P*=0.028) and end-systolic (21ul vs. 14ul, *P*<0.001) volumes compared with controls (Figure 2A).

**Figure 2.**
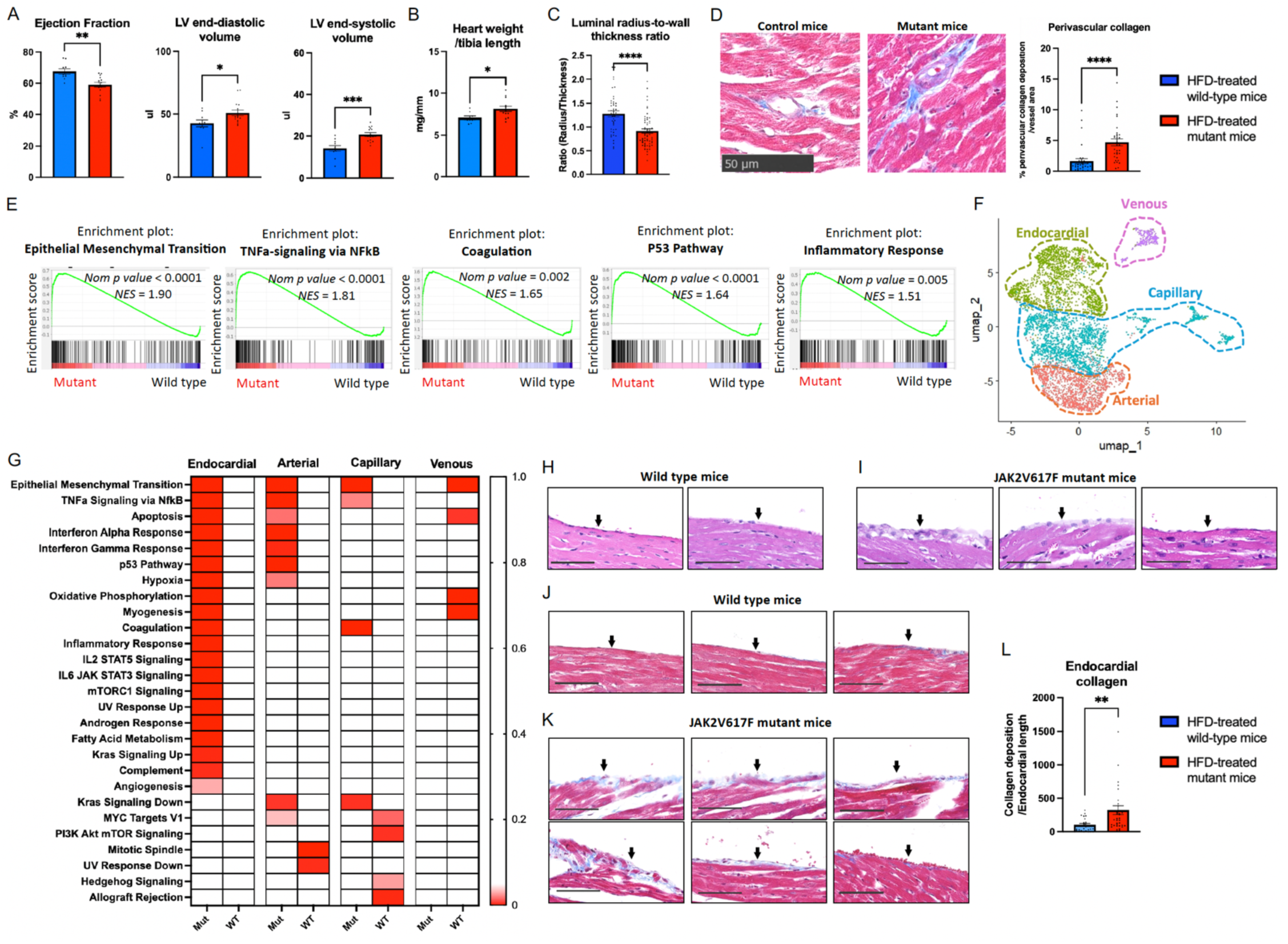
JAK2V617F Mutant hematopoiesis Promotes a Distinct Cardiac Microvascular Disease Phenotype When under a High-Fat Diet Challenge. (**A**) Transthoracic echocardiography showing reduced LV ejection fraction and increased LV end-diastolic volume and end-systolic volumes in HFD-treated JAK2V617F mutant mice (red) compared with wild-type controls (blue) (n=10-15 mice per group from 3 independent experiments). (**B**) Heart weight normalized to tibia length in JAK2V617F mutant and wild-type control mice after 5 weeks of HFD treatment (n=9-13 mice per group; 3 independent experiments). (**C**) Quantification of the luminal radius-to-wall thickness ratio, measured in nonoverlapping coronary arterioles from JAK2V617F mutant mice and wild-type mice (6-8 mice per group; 72-100 microvessels analyzed per group). (**D**) Representative Masson’s trichrome staining (left) and quantitation (right) of perivascular collagen deposition around coronary arterioles in JAK2V617F mutant and wild-type mice (3 mice per group; 33 microvessels analyzed per group). Each dot represents an individual microvessel. (**E**) GSEA of differentially expressed genes in cardiac ECs from HFD-treated mice with JAK2V617F mutant versus wild-type blood cells. (**F**) UMAP visualization of cardiac EC subpopulations. (**G**) GSEA of endocardial, arterial, venous, and capillary ECs from HFD-treated JAK2V617F mutant (Mut) versus wild-type (WT) mice. The color scale on the right indicates false discovery rate (FDR) values. (**H-K**) Representative H&E (H-I) and Masson’s trichrome (J-K) staining of the endocardium (arrows) in HFD-treated wild-type and JAK2V617F mutant mice. Scale bar: 50um. (**L**) Quantitation of subendocardial collagen deposition across nonoverlapping endocardium segments from JAK2V617F mutant and wild-type mice (3 mice per group; 31-36 segments analyzed per group, average segment length 62um). \**P*<0.05, \*\**P*<0.01, \*\*\**P*<0.001, \*\*\*\**P*<0.0001 Data are presented as mean ± SEM and *P* value was calculated using an unpaired two-tailed Student’s t-test.

Pathological evaluation revealed an increased heart weight-to-tibia length ratio in mutant mice (Figure 2B). Mutant mice fed an HFD exhibited marked coronary arteriole stenosis (luminal radius-to-wall thickness ratio 0.9 vs. 1.3, *P*<0.0001) (Figure 2C) and significantly increased perivascular collagen deposition (2.9-fold, *P*<0.0001) (Figure 2D). No gross atherosclerosis, epicardial coronary stenosis, or myocardium infarction was detected.

To investigate how HFD affects endothelial function in the presence of JAK2V617F-mutant blood cells, we performed scRNAseq on unfractionated heart cells from mutant and wild-type mice after diet treatment. Cardiac ECs from HFD-treated mutant mice showed enrichment of EndMT, TNFa/NFkB signaling, coagulation, P53, and inflammatory response pathways, indicating a shift toward a dysfunctional, pro-inflammatory, and mesenchymal/fibrotic state (Figure 2E). This endothelial reprogramming could explain the observed phenotype of impaired cardiac function, coronary arteriole stenosis, and perivascular fibrosis.

To further define the cellular players in JAK2V617F mutant blood cell-induced endothelial dysfunction, we performed an integrated cluster analysis of the EC population and identified four distinct EC populations: arterial ECs (Sema3g, Edn1, Nos3, Fbln5), venous ECs (Nr2f2, Lyve1, Igfbp5), capillary ECs (Rgcc, Aqp1, Cd36, Gpihbp1), and endocardial ECs (Nfatc, Nrg1, Tbx20, Cdh11)^48,49^ (Figure 2F). The most pronounced transcriptomic changes occurred in endocardial ECs from HFD-treated mutant mice, which showed robust upregulation of pathways involved in inflammation (e.g., TNFa signaling, IFNα/γ signaling, inflammatory response), metabolic reprogramming (e.g., MTORC1 signaling, fatty acid metabolism, oxidative phosphorylation), stress responses (e.g., P53 pathway, apoptosis, hypoxia), and EndMT (Figure 2G). In line with these findings, histological evaluation revealed endocardial disruption — characterized by irregular lining, prominent nuclei, and cellular enlargement or thickening — in 7 of 8 mutant mice but in none of the 6 wild-type controls examined (Figure 2H-I). Masson’ trichrome staining showed significantly increased subendocardium collagen deposition in mutant mice (3.2-fold, *P*=0.002) (Figure 2J-L). Taken together, these findings indicate that JAK2V617F-mutant blood cells combined with HFD induced a distinctive CVD phenotype characterized by cardiac microvascular dysfunction, including coronary arteriole stenosis, perivascular fibrosis, and increased endothelial inflammation. LV mass is increased, whereas LVEF remains relatively preserved. Notably, an unexpected yet consistent finding is disruption of the endocardial endothelium.

The endocardium is the innermost layer of the heart chambers and is composed of a thin monolayer of specialized ECs that line the cardiac interior. Endocardial cells are among the earliest endothelial populations to emerge during heart development and contribute to coronary vasculature formation during both embryogenesis^50^ and postnatal life^51^. Given their critical role in maintaining cardiac homeostasis and promoting repair and regeneration after cardiac injury^52,53^, the impact of mutant blood cells on endocardial EC function warrants further investigation.

### MPL is Predominantly Expressed in a Subset of Endocardial ECs

The TPO receptor MPL is expressed on various vascular ECs^11–14^, and TPO/MPL signaling has been implicated in cardiovascular regulation^12,13,15–17^. While MPL expression has been reported in human umbilical cord ECs^11,12^, mouse liver^13^ and pulmonary^14^ ECs, MPL expression within the heart has not been characterized. To address this, we examined MPL expression in murine cardiac ECs using multiple complementary approaches.

First, RT-PCR was performed on ECs (CD45^-^CD31^+^) isolated from heart, lung, marrow, and spleen of a wild-type mouse. cDNA generated by reverse transcription was amplified using primers targeting a 111-bp sequence spanning mouse MPL exons 2-3 (Mm00440306_g1, ThermoFisher®). MPL transcripts were consistently detected in ECs from all tissues (Figure 3A). Similarly, using a human MPL-specific primer, we detected MPL transcripts in human umbilical vein ECs (HUVECs), consistent with previous reports^11,12^ (data not shown).

**Figure 3.**
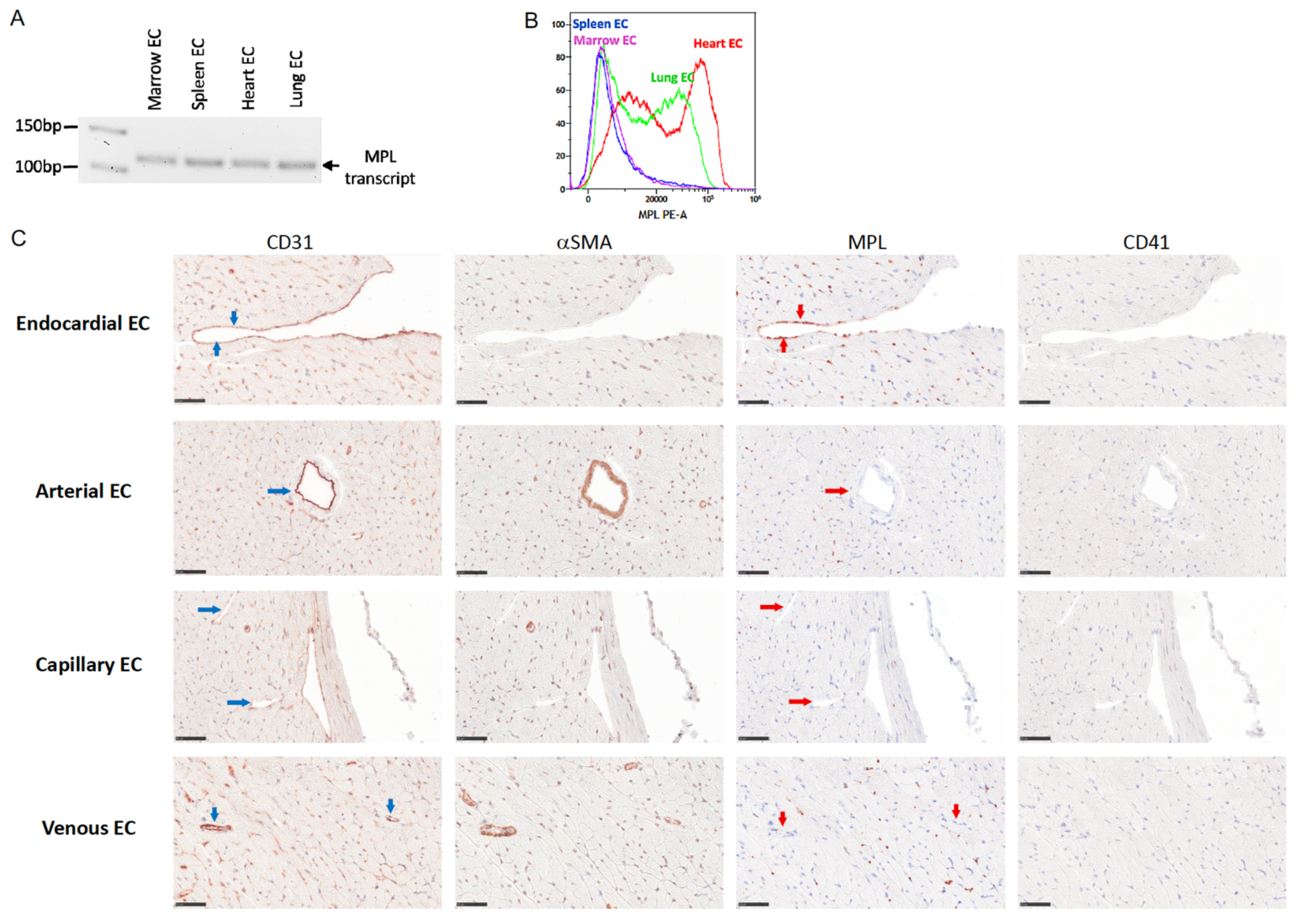
MPL is Selectively Expressed in a subset of Endocardial ECs. (**A**) MPL gene expression in isolated vascular ECs (CD45^-^CD31^+^) from heart, lung, bone marrow, and spleen of wild-type C57B/L6 mice, measured by RT-PCR. (**B**) Cell surface MPL protein expression in ECs from heart, lung, bone marrow, and spleen, assessed by flow cytometry using the anti-MPL antibody AMM2. (**C**) Representative immunohistochemical staining of mouse heart showing MPL localization within distinct endothelial subtypes. Blue arrows indicate vascular ECs with positive CD31 staining; red arrows denote MPL staining in the same cells, demonstrating positive expression in some endocardial ECs but not in arterial, capillary, venous ECs. Scale bar, 50um; magnification, 40x.

Second, cell surface MPL protein expression was evaluated by flow cytometry analysis using the AMM2 antibody, a rat anti-mouse MPL neutralizing antibody (Cat. 10401, IBL-America®). In line with our prior findings in MPL-GFP reporter mice (where GFP marks MPL transcriptional activity)^22,33^, MPL expression was highest in cardiac ECs, followed by lung ECs, and comparatively lower in marrow and spleen ECs (Figure 3B).

Third, we performed immunohistochemistry with a rabbit anti-mouse MPL antibody (ab232755, Abcam®) to localize MPL expression within the heart’s endothelial subtypes — endocardial, arterial, venous, and capillary. CD31 (or Pecam1) was used to identify vascular ECs, and CD41 to exclude platelets which are MPL-positive. The endocardium was identified by its specific location lining the cardiac chambers and papillary muscles, whereas arteries were recognized by its thick media containing abundant concentric smooth muscle layers, veins by their thin wall with sparse smooth muscle, and capillaries by their single endothelial layer (no smooth muscle)^40^. Strikingly, MPL expression was observed predominantly in a subset of endocardial ECs, whereas arterial, venous, and capillary ECs were largely negative (Figure 3C).

Previously, we showed that MPL expression is upregulated in JAK2V617F-mutant ECs compared with wild-type ECs^22,27,28^, and that genetic ablation of endothelial MPL suppresses JAK2V617F-induced EndMT and prevents CVD development in mice with JAK2V617F-mutant ECs^22^. Building on our new finding that MPL is selectively expressed in endocardial ECs, a subpopulation with critical roles in maintaining cardiac function and mediating repair and regeneration following cardiac injury^52,53^, and that these cells exhibit marked transcriptomic and histological perturbations in mice with JAK2V617F-mutant hematopoiesis under metabolic stress (Figure 2), we next investigated whether pharmacological inhibition of TPO/MPL signaling using the MPL-neutralizing antibody AMM2 could prevent CVD development *in vivo* in mice bearing JAK2V617F-mutant blood cells.

### AMM2 Prevents CVD Development in Mice with JAK2V617F Mutant Blood Cells by Restoring Endocardial Integrity and Microvascular function

AMM2 was first described by Yoshihara et al. in 2007 as an anti-MPL neutralizing antibody, where a short course (6 days, 1mg/kg) reduced cell-cycle regulators and increased Lin^-^cKit^+^Sca1^+^ (or LSK) hematopoietic stem/progenitor cell cycling and engraftment *in vivo*^29^. Olson et al. subsequently reported that AMM2 (1mg/kg) administered at the time of irradiation did not reduce total marrow megakaryocytes, though it impaired their migration to the endosteal niche^30^. Li et al. demonstrated that AMM2 treatment (0.1mg/kg per day for 4 days) enhanced the therapeutic effect of chemotherapy (Ara-C) in a leukemia murine model, likely by promoting leukemia stem cell cycling^31^. In addition, AMM2 has been widely used as an MPL-specific antibody to measure receptor expression by flow cytometry and Western blot^32–34^.

Since TPO/MPL signaling is a key regulator of HSC function^9^, we first assessed whether AMM2 treatment would impair hematopoiesis. In a pilot study, wild-type mice received a single intravenous injection of AMM2 at 0.1mg/kg. Peripheral blood platelet counts remained stable throughout a 14-day follow-up period (Figure 4A). We then treated wild-type mice with 0.1mg/kg AMM2 intravenously every 10 days for a total of four doses to evaluate potential hematopoietic effects during extended treatment. No significant differences were observed in peripheral blood counts or in bone marrow megakaryocytes and Lin^-^cKit^+^Sca1^+^ (LSK) hematopoietic stem and progenitor cells between AMM2- and PBS-treated mice (Figure 4B). Therefore, the 0.1mg/kg AMM2 regimen was used for all subsequent *in vivo* experiments as a haematopoietically safe dose.

**Figure 4.**
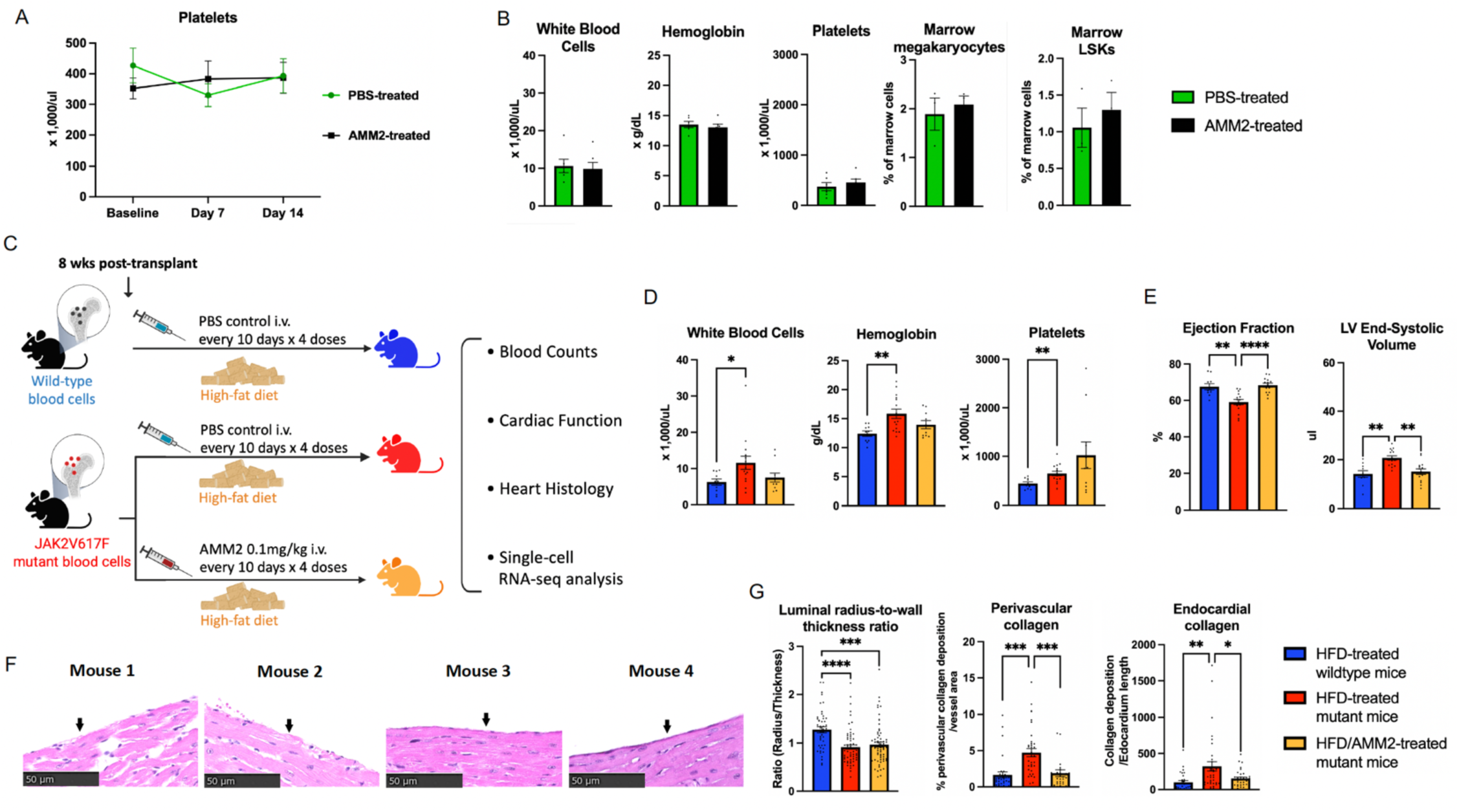
AMM2 Prevents CVD Development in Mice with JAK2V617F Mutant Blood Cells even under High-Fat Diet Challenge. (**A**) Peripheral blood platelet counts in wild-type mice following a single dose of saline (PBS) or AMM2 (n=4-7 mice per group). (**B**) Peripheral blood counts (n=6 per group) and bone marrow megakaryocyte and LSK hematopoietic stem and progenitor cells (n=3 mice per group) in wild-type mice after four doses of PBS or AMM2. (**C**) Schematic illustration of the experimental design showing the HFD feeding and AMM2 treatment schedule. (**D-E**) Peripheral blood counts (D) and LV ejection fraction and volumes (E) in wild-type mice on HFD, JAK2V617F mutant mice on HFD, and JAK2V617F mutant mice on HFD+AMM2 (n=10-15 mice per group from 3 independent experiments). (**F**) Representative H&E staining of the endocardium (arrows) from four JAK2V617F mutant mice treated with both HFD and AMM2. (**G**) Quantification of (left) coronary arteriole luminal radius-to-wall thickness ratio (8 mice per group; a total of 84-100 microvessels analyzed per group), (middle) coronary arteriole perivascular collagen deposition (3 mice per group; a total of 33 microvessels analyzed per group), and (right) subendocardial collagen deposition (3 mice per group; a total of 31-36 segments analyzed per group, average segment length 62um) in wild-type mice on HFD, JAK2V617F mutant mice on HFD, and JAK2V617F mutant mice on HFD+AMM2. \**P*<0.05, \*\**P*<0.01, \*\*\**P*<0.001, \*\*\*\**P*<0.0001 Data are presented as mean ± SEM and *P* value was calculated using an unpaired two-tailed Student’s t-test (panels A and B) or one-way ANOVA (panels D, E, and G)

We generated chimeric mice harboring JAK2V617F mutant blood cells and wild-type ECs through bone marrow transplantation. By 8 weeks post-transplantation, JAK2V617F mutant mice had already developed a classic MPN phenotype, including leukocytosis, erythrocytosis, and thrombocytosis, compared to wild-type controls (as shown in Figure 1B)^21^. At this time, JAK2V617F mutant mice were assigned to one of two groups for a total of five weeks of treatment: (1) HFD alone or (2) HFD plus AMM2, administered intravenously at 0.1mg/kg every 10 days for a total of four doses. Chimeric mice with wild-type blood cells and ECs served as controls and received HFD alone (Figure 4C). Consistent with findings in wild-type mice, AMM2 treatment did not alter peripheral blood counts in JAK2V617F mutant mice (Figure 4D) but provided clear cardiovascular protection: whereas mutant mice on HFD alone developed cardiac dysfunction (see Figure 2A), AMM2-treated mice maintained normal LV ejection fraction and LV volumes comparable to wild-type controls (Figure 4E). The experiment was repeated three times, each yielding consistent results. To ensure that the cardioprotective effect was not attributable to the antibody solvent (1% bovine serum albumin [BSA] and 0.05% sodium azide [NaN_3_]), we repeated the experiment using a customized AMM2 formulation without BSA and NaN_3_ (IBL-America®) and observed identical results (data not shown). Histological analysis revealed that, compared to HFD treatment alone (Figure 2I and 2K), AMM2 markedly restored endocardial integrity in all HFD-treated JAK2V617F mutant mice examined (n=6) (Figure 4F). Quantitative assessment demonstrated persistent coronary arteriole stenosis; however, perivascular collagen fibrosis and subendocardium collagen deposition were significantly reduced in AMM2-treated mice (Figure 4G), suggesting that AMM2 improved both cardiac microvascular function and endocardial integrity.

To investigate the molecular mechanisms by which AMM2 prevents HFD-induced cardiovascular dysfunction in mice with JAK2V617F-mutant blood cells, we performed scRNAseq on unfractionated heart cells from mutant mice on HFD and from mutant mice on HFD plus AMM2. AMM2 treatment markedly suppressed inflammatory (e.g., TNFα/NFκB signaling, IFNα/γ signaling) and stress response (e.g., UV response, apoptosis, P53, hypoxia) pathways when all cardiac ECs were analyzed collectively (Figure 5A). Among the individual endothelial subtypes, endocardial ECs displayed the most pronounced transcriptomic changes (Figure 5B), which aligns with our finding that MPL expression is largely restricted to endocardial ECs (Figure 3C). These molecular changes, combined with the histological findings of enhanced endocardial integrity and reduced endocardium and perivascular fibrosis in AMM2-treated mice (Figure 4F-G), indicate that AMM2 treatment promotes a protective endothelial phenotype and minimizes cell death and maladaptive remodeling in HFD-treated JAK2V617F mutant mice.

**Figure 5.**
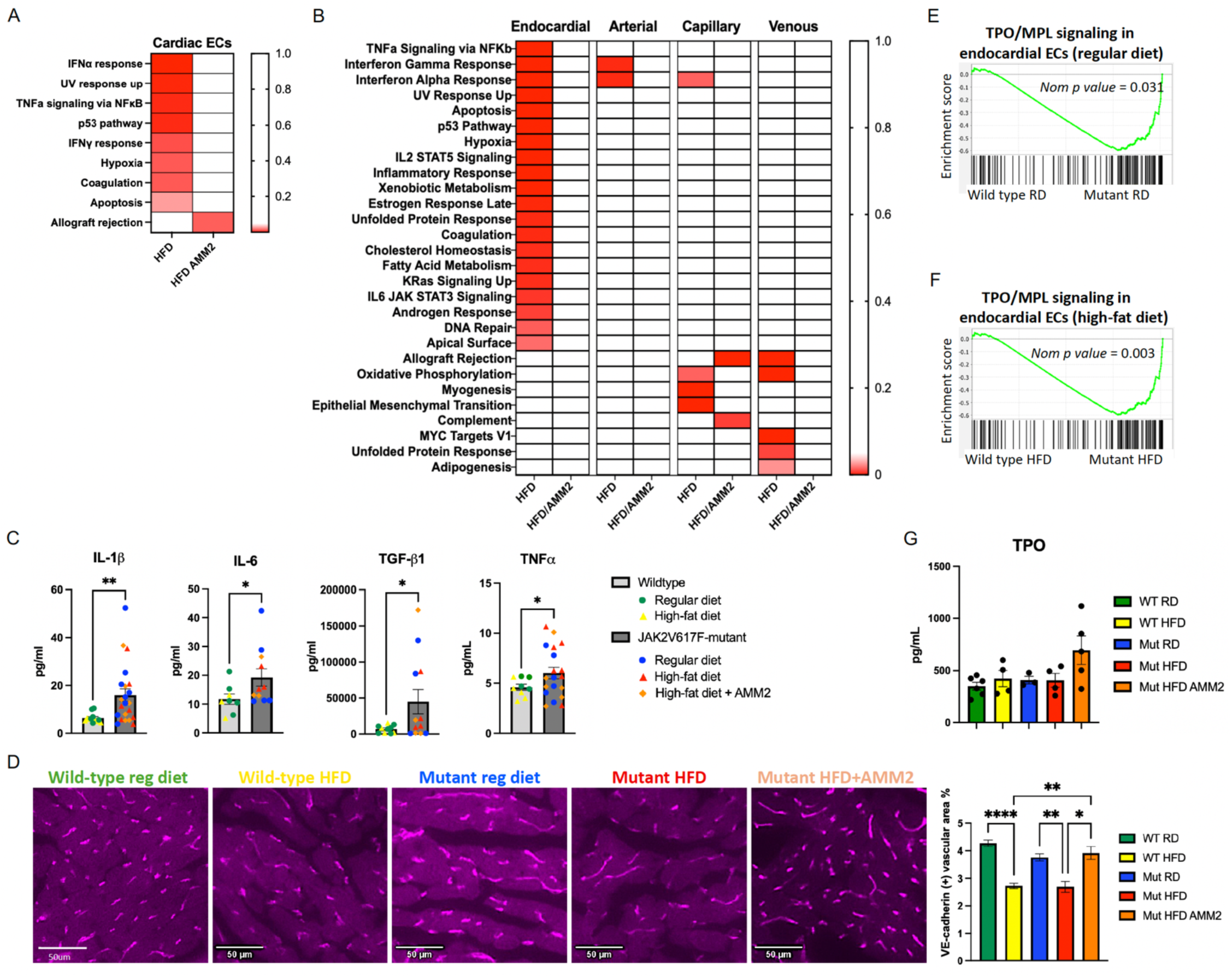
AMM2 Treatment Improves Cardiac Microvascular Function in Mice with JAK2V617F-mutant Hematopoiesis Despite Persistent Systemic Inflammation. (**A-B**) GSEA of differentially expressed genes in all cardiac ECs (A) and EC subpopulations (B) from JAK2V617F mutant mice treated with HFD vs. HFD+AMM2. (**C**) Plasma levels of IL1β (n=12-22 mice per group), IL6 (n=6-9 mice per group), TGFβ (n=10-12 mice per group), TNFα (n=9-12 mice per group) were significantly elevated in JAK2V617F mutant mice compared to wild-type controls. (**D**) Representative 20x images and quantification of VE-cadherin^+^ vasculature (magenta) in the hearts of wild-type (WT) and JAK2V617F mutant (Mut) mice after 5 weeks of regular diet (RD), HFD, or HFD plus AMM2 treatment, demonstrating increased microvascular density in mutant mice treated with HFD+AMM2 compared with HFD alone (n=2 mice per group, with 19-30 20x image fields analyzed per group). (**E-F**) TPO/MPL signaling genes were markedly upregulated in endocardial ECs from JAK2V617F mutant mice compared with wild-type mice under both regular diet (E) and HFD (F) conditions. (**G**) Plasma TPO levels across experimental groups (n=3-5 mice per group). * *P*<0.05, ** *P*<0.01, **** *P*<0.0001. Data are presented as mean ± SEM and *P* value was calculated using an unpaired two-tailed Student’s t-test (panel C) or one-way ANOVA (panels D and G).

Consistent with chronic inflammation as a hallmark of JAK2V617F-mutant hematopoiesis^54^, interleukin-1 beta (IL1β), interleukin-6 (IL6), transforming growth factor beta 1 (TGFβ), tumor necrosis factor-alpha (TNFα) were significantly elevated in JAK2V617F mutant mice compared to wild-type mice (Figure 5C); however, their levels were not affected by dietary intervention or AMM2 treatment (Supplementary Figure 1). Therefore, while the JAK2V617F mutant hematopoiesis induced persistent systemic inflammation with elevated IL-1β, IL-6, TGFβ, and TNFα, AMM2 was able to suppress inflammation in cardiac ECs, particularly within the endocardial EC population. Such attenuation of endothelial inflammation may shift the coronary microvasculature from a dysfunctional to a regenerative state by enhancing endothelial cell proliferation^55^. Supporting this model, *in vivo* VE-cadherin labeling revealed that, while HFD reduced coronary microvascular density in both wild-type and JAK2V617F mutant mice, AMM2 treatment was able to increase coronary microvasculature in mutant mice (Figure 5D).

To investigate how AMM2 selectively targets endocardial ECs, we compiled a list of 130 TPO/MPL signaling-related genes by integrating multiple sources, including (1) the TPO Signaling pathway from GeneGlobe (QIAGEN®), (2) the BioCarta TPO pathway gene set, and (3) Bhat et al^56^. We then assessed pathway enrichment across various cardiac EC subpopulations using our scRNAseq dataset. Notably, endocardial ECs — but not arterial, capillary, or venous ECs — from mice carrying JAK2V617F-mutant blood cells exhibited significant enrichment of TPO/MPL signaling genes compared with wild-type controls under both regular diet (Figure 5E) and high-fat diet (Figure 5F) conditions. Such changes were not observed in arterial, capillary, or venous cardiac ECs (Supplementary Figure 2). TPO/MPL signaling is primarily regulated by ligand-dependent activation through TPO binding. Although we did not detect significant differences in peripheral blood TPO levels among experimental groups (Figure 5G), these measurements may not reflect local cytokine dynamics within the cardiac microenvironment. Therefore it is plausible that chronic inflammation associated with JAK2V617F-mutant hematopoiesis, characterized by elevated cytokines such as IL-6^57^, enhances local TPO production or bioavailability within vascular ECs^13^. Such paracrine or autocrine signaling could preferentially activate MPL in endocardial ECs, thereby driving the observed transcriptional program.

Together, these results identify endocardial ECs as an important target of JAK2V617F mutant hematopoiesis and demonstrate that MPL is selectively expressed in this endothelial subset. The anti-MPL neutralizing antibody AMM2 alleviates stress (e.g., HFD)-induced endocardial injury and cardiovascular dysfunction in JAK2V617F mutant mice, likely through suppression of endocardial inflammation and enhancement of coronary vascular repair and/or regeneration.

## Discussion

In this study, we show that mice harboring JAK2V617F-mutant blood cells develop a distinctive form of cardiac microvascular disease, both at baseline and after a high-fat/high-cholesterol diet challenge, characterized by increased LV mass with preserved LVEF. Under HFD stress, a modest reduction in LVEF emerges, accompanied by clear evidence of endocardial endothelial dysfunction. Histologically, this phenotype is marked by coronary arteriole stenosis, perivascular fibrosis, endocardial disruption, and reduced microvascular density — hallmarks of cardiac microvascular dysfunction — in the absence of overt epicardial coronary artery disease.

Unlike prior studies using hyperlipidemic *Ldlr^-/-^* mice^58^ or invasive surgical models (e.g., coronary ligation, transverse aorta constriction)^59^ to link JAK2V617F-mutant blood cells to atherosclerosis and inflammation, our study employs a noninvasive high-fat/high-cholesterol Western diet model that represents a common and modifiable cardiovascular risk factor in developed countries^60^. Western-style diet is a well-established preventable risk factor for CVD, and even a single high-fat meal can acutely impair endothelial function^61,62^. Using this clinically relevant model, we show that JAK2V617F-mutant blood cells alone are sufficient to induce distinctive cardiac microvascular disease both at baseline and following a high-fat/high-cholesterol diet challenge (Figures 1-2). In the adult heart, a tightly regulated interplay among ECs (including endocardial ECs), fibroblasts, and immune cells preserves microenvironmental homeostasis, enabling adaptation to stress and initiation of reparative responses^63^. Our findings reveal that mutant hematopoiesis induces microvascular injury and triggers adaptive remodeling of the cardiac microenvironment, preserving LV systolic function via heightened endothelial inflammatory and stress signaling. Under metabolic stress (HFD), this compensatory response amplifies and increasingly depends on endocardial EC engagement. Prolonged HFD exposure ultimately disrupts this balance, leading to endocardial endothelial exhaustion, fibroblast activation, and maladaptive remodeling characterized by coronary arteriole stenosis, endocardial dysfunction, perivascular fibrosis, and increased LV volume/mass. These features define a pathophysiologic continuum leading to overt microvascular dysfunction in JAK2V617F-mutant mice under metabolic challenge. Unlike our prior model which demonstrated overt heart failure with thrombosis, cardiomyopathy, and both macro- and micro-vasculopathy driven by mutant blood cells and mutant vascular ECs^21^, this study offers a novel, noninvasive model of coronary microvascular disease.

Previous studies have linked JAK2V617F-mutant blood cells to vascular pathology: Poisson et al. showed mutant erythrocytes induce endothelial oxidative stress and arterial vasoconstriction in the aorta and femoral arteries^64^; Kimishima et al. reported worsened hypoxia-induced pulmonary hypertension with pulmonary arterial wall thickening^65^; and Molinaro et al. demonstrated exacerbated endothelial damage and arterial thrombosis following carotid artery denudation^66^. In contrast, our study focuses on cardiac ECs and provides several novel insights. We demonstrated for the first time that the TPO receptor MPL is selectively expressed in murine endocardial ECs, a population critical for coronary vasculature formation, cardiac repair, and regeneration^52,53^. Single-cell RNA-seq identified these cells as the primary target of mutant hematopoiesis-driven CVD under cardiometabolic stress, showing activation of inflammatory, stress-response, and EndMT programs plus endocardial disruption. Importantly, the MPL-neutralizing antibody AMM2 ameliorated high-fat/high-cholesterol diet-induced abnormalities by restoring endocardial integrity, increasing microvascular density, and downregulating inflammatory/stress-response signatures in endocardial ECs. Notably, low-dose AMM2 achieved these cardioprotective effects without altering peripheral blood counts in wild-type mice. Given the established role of MPL in MPN pathogenesis^67–73^, our findings position endocardial ECs as a novel mediator of mutant hematopoiesis–driven cardiovascular injury and highlight cardiac endothelial MPL as a potential therapeutic target for preventing CVD in JAK2V617F-positive MPN or clonal hematopoiesis.

During embryonic development, endocardial ECs are essential for cardiac valves and septa formation through EndMT^74^. They also contribute to coronary vasculature development embryonically^50,75,76^ and postnatally^51,77–80^ in zebrafish and mice, while supporting cardiac function, repair, and regeneration after injury^52,53^. In our study, AMM2 treatment suppressed inflammatory and stress-response pathways in endocardial ECs, alleviated endocardial injury, increased coronary microvascular density, and improved cardiac function in HFD-treated JAK2V617F mutant mice, despite persistent systemic inflammation (elevated IL-1β, IL-6, TGFβ and TNFα; Figures 4-5). These findings suggest that endocardial EC-mediated coronary vascular repair/regeneration is a central mechanism of AMM2’s cardioprotection. Although clonal hematopoiesis and associated systemic inflammation may be inevitable with aging, enhancing the intrinsic repair capacity of tissue-resident ECs offers a promising strategy to prevent aging-related cardiovascular diseases linked to clonal hematopoiesis. Further studies are required to understand the role of endocardial TPO/MPL signaling in JAK2V617F-induced cardiac microvascular disease and whether AMM2 acts directly on endocardial TPO/MPL signaling or indirectly via hematopoietic cells.

Our model may have clinical application by recapitulating key pathological features of non-obstructive coronary arteries and heart failure with preserved ejection fraction (HFpEF), in which coronary microvascular disease is increasingly recognized as a central pathogenic mechanism^81,82^. Consistent with this framework, CHIP has been linked to increased prevalence of coronary microvascular dysfunction in patients with early, non-obstructive coronary artery disease^83^. Large observational studies demonstrate that the association between CHIP and incident heart failure is independent of epicardial coronary artery disease, implicating non-ischemic, microvascular, or myocardial mechanisms^84^. In a recent case–control study of incident heart failure, CHIP was associated with higher natriuretic peptide levels, with subgroup analyses indicating a stronger link to HFpEF — particularly in individuals under 65 years — than to heart failure with reduced ejection fraction^85^. Moreover, in a large two-cohort study, TET2-associated CHIP conferred more than a twofold increased risk of HFpEF, independent of traditional cardiovascular risk factors and coronary artery disease^86^. Our model has provided an ideal platform for testing dietary/metabolic interventions and other therapeutic options such as anti-inflammatory agent to treat or prevent cardiac microvascular disease in clonal hematopoiesis. Although congestive heart failure and diastolic function were not specifically assessed in our murine model, it is important to note that conventional echocardiographic indices of diastolic function lack sensitivity for the diagnosis of HFpEF^87^.

In summary, our study has established a model in which JAK2V617F-mutant hematopoiesis drives cardiac microvascular injury and maladaptive remodeling, consistent with clinical observations in patients with clonal hematopoiesis. This process is amplified by metabolic stress and mediated, at least in part, through selective vulnerability of endocardial ECs. We discovered MPL expression in endocardial ECs and showed that pharmacologic MPL inhibition with AMM2 attenuates endocardial inflammation, preserves integrity, and increases coronary vascular density, and improves cardiac function. These findings mechanistically link mutant hematopoiesis to human microvascular heart disease and highlight TPO/MPL signaling as a potential targetable pathway to reduce cardiovascular risk in JAK2V617F-positive CHIP or MPNs.

## ACKNOWLEDGEMENTS

This work was supported by the VA Merit Award BX003947 and BX005584 (H.Z.) and NIH R01 CA266294 (H.Z.). We thank Ms. Isabel Zheng (Oldfield Middle School, Greenlawn, NY) for her contribution to the graphic abstract.

## AUTHOR CONTRIBUTIONS

X.Y. performed all experiments, analyzed the data, and prepared the figures; K.M. contributed to the transgenic murine model studies and scRNAseq data analysis; X.S. conducted histology and plasma sample analysis and quantified VE-cadherin-positive vascular areas; F.Z. analyzed perivascular fibrosis and subendocardial collagen deposition; K.Z. performed blinded quantification of VE-cadherin-positive vascular area; H. Zheng provided scientific and technical expertise in cardiovascular functional assessments and histologic evaluations, interpreted the results, and drafted the manuscript; H. Zhan conceived and supervised the study, analyzed data, interpreted results, and drafted the manuscript. All authors reviewed and approved the manuscript.

## REFERENCES

1. Landolfi R, Di Gennaro L, Falanga A. Thrombosis in myeloproliferative disorders: pathogenetic facts and speculation. Leukemia. 2008;22:2020–2028. doi: 10.1038/leu.2008.253

2. Bick AG, Weinstock JS, Nandakumar SK, Fulco CP, Bao EL, Zekavat SM, Szeto MD, Liao X, Leventhal MJ, Nasser J, et al. Inherited causes of clonal haematopoiesis in 97,691 whole genomes. Nature. 2020;586:763–768. doi: 10.1038/s41586-020-2819-2

3. Pershad Y, Uddin MM, Xue L, Haessler J, Collins JM, Mack TM, Glick E, Glaser V, Zhao K, Jaiswal S, et al. Correlates and consequences of clonal hematopoiesis expansion rate: a 16-year longitudinal study of 6976 women. Blood. 2025;146:1078–1087. doi: 10.1182/blood.2025028417

4. Jaiswal S, Fontanillas P, Flannick J, Manning A, Grauman PV, Mar BG, Lindsley RC, Mermel CH, Burtt N, Chavez A, et al. Age-related clonal hematopoiesis associated with adverse outcomes. N Engl J Med. 2014;371:2488–2498. doi: 10.1056/NEJMoa1408617

5. Jaiswal S, Natarajan P, Silver AJ, Gibson CJ, Bick AG, Shvartz E, McConkey M, Gupta N, Gabriel S, Ardissino D, et al. Clonal Hematopoiesis and Risk of Atherosclerotic Cardiovascular Disease. N Engl J Med. 2017;377:111–121. doi: 10.1056/NEJMoa1701719

6. Yokokawa T, Misaka T, Kimishima Y, Wada K, Minakawa K, Kaneshiro T, Yoshihisa A, Ikeda K, Takeishi Y. Clonal Hematopoiesis and JAK2V617F Mutations in Patients With Cardiovascular Disease. JACC CardioOncol. 2021;3:134–136. doi: 10.1016/j.jaccao.2021.01.001

7. Flynn S, Schuermans A, Uddin MM, Nakao T, Viscosi V, Libby P, Natarajan P, Honigberg MC. Clonal Hematopoiesis and Incident Heart Failure. JAMA Cardiol. 2025. doi: 10.1001/jamacardio.2025.4603

8. Wu J, Zhang L, Vaze A, Lin S, Juhaeri J. Risk of Wernicke’s encephalopathy and cardiac disorders in patients with myeloproliferative neoplasm. Cancer Epidemiol. 2015;39:242–249. doi: 10.1016/j.canep.2015.01.014

9. Kaushansky K. Thrombopoietin, the Primary Regulator of Platelet Production: From Mythos to Logos, a Thirty-Year Journey. Biomolecules. 2024;14. doi: 10.3390/biom14040489

10. Varghese LN, Defour JP, Pecquet C, Constantinescu SN. The Thrombopoietin Receptor: Structural Basis of Traffic and Activation by Ligand, Mutations, Agonists, and Mutated Calreticulin. Front Endocrinol (Lausanne). 2017;8:59. doi: 10.3389/fendo.2017.00059

11. Methia N, Louache F, Vainchenker W, Wendling F. Oligodeoxynucleotides antisense to the proto-oncogene c-mpl specifically inhibit in vitro megakaryocytopoiesis. Blood. 1993;82:1395–1401.

12. Brizzi MF, Battaglia E, Montrucchio G, Dentelli P, Del Sorbo L, Garbarino G, Pegoraro L, Camussi G. Thrombopoietin stimulates endothelial cell motility and neoangiogenesis by a platelet-activating factor-dependent mechanism. Circ Res. 1999;84:785–796. doi: 10.1161/01.res.84.7.785

13. Cardier JE, Dempsey J. Thrombopoietin and its receptor, c-mpl, are constitutively expressed by mouse liver endothelial cells: evidence of thrombopoietin as a growth factor for liver endothelial cells. Blood. 1998;91:923–929.

14. English J, Dhanikonda S, Tanaka KE, Koba W, Eichenbaum G, Yang WL, Guha C. Thrombopoietin mimetic reduces mouse lung inflammation and fibrosis after radiation by attenuating activated endothelial phenotypes. JCI Insight. 2024;9. doi: 10.1172/jci.insight.181330

15. Eguchi M, Masuda H, Kwon S, Shirakura K, Shizuno T, Ito R, Kobori M, Asahara T. Lesion-targeted thrombopoietin potentiates vasculogenesis by enhancing motility and enlivenment of transplanted endothelial progenitor cells via activation of Akt/mTOR/p70S6kinase signaling pathway. J Mol Cell Cardiol. 2008;45:661–669. doi: 10.1016/j.yjmcc.2008.08.002

16. Lupia E, Spatola T, Cuccurullo A, Bosco O, Mariano F, Pucci A, Ramella R, Alloatti G, Montrucchio G. Thrombopoietin modulates cardiac contractility in vitro and contributes to myocardial depressing activity of septic shock serum. Basic Res Cardiol. 2010;105:609–620. doi: 10.1007/s00395-010-0103-6

17. Li K, Sung RY, Huang WZ, Yang M, Pong NH, Lee SM, Chan WY, Zhao H, To MY, Fok TF, et al. Thrombopoietin protects against in vitro and in vivo cardiotoxicity induced by doxorubicin. Circulation. 2006;113:2211–2220. doi: 10.1161/CIRCULATIONAHA.105.560250

18. Sozer S, Fiel MI, Schiano T, Xu M, Mascarenhas J, Hoffman R. The presence of JAK2V617F mutation in the liver endothelial cells of patients with Budd-Chiari syndrome. Blood. 2009;113:5246–5249. doi: 10.1182/blood-2008-11-191544

19. Rosti V, Villani L, Riboni R, Poletto V, Bonetti E, Tozzi L, Bergamaschi G, Catarsi P, Dallera E, Novara F, et al. Spleen endothelial cells from patients with myelofibrosis harbor the JAK2V617F mutation. Blood. 2013;121:360–368. doi: 10.1182/blood-2012-01-404889

20. Helman R, Pereira WO, Marti LC, Campregher PV, Puga RD, Hamerschlak N, Chiattone CS, Santos FPS. Granulocyte whole exome sequencing and endothelial JAK2V617F in patients with JAK2V617F positive Budd-Chiari Syndrome without myeloproliferative neoplasm. Br J Haematol. 2018;180:443–445. doi: 10.1111/bjh.14327

21. Castiglione M, Jiang YP, Mazzeo C, Lee S, Chen JS, Kaushansky K, Yin W, Lin RZ, Zheng H, Zhan H. Endothelial JAK2V617F mutation leads to thrombosis, vasculopathy, and cardiomyopathy in a murine model of myeloproliferative neoplasm. J Thromb Haemost. 2020. doi: 10.1111/jth.15095

22. Zhang H, Kafeiti N, Masarik K, Lee S, Yang X, Zheng H, Zhan H. Decoding Endothelial MPL and JAK2V617F Mutation: Insight Into Cardiovascular Dysfunction in Myeloproliferative Neoplasms. Arterioscler Thromb Vasc Biol. 2024;44:1960–1974. doi: 10.1161/ATVBAHA.124.321008

23. Bischoff J. Endothelial-to-Mesenchymal Transition. Circ Res. 2019;124:1163–1165. doi: 10.1161/CIRCRESAHA.119.314813

24. Cho JG, Lee A, Chang W, Lee MS, Kim J. Endothelial to Mesenchymal Transition Represents a Key Link in the Interaction between Inflammation and Endothelial Dysfunction. Front Immunol. 2018;9:294. doi: 10.3389/fimmu.2018.00294

25. Dejana E, Hirschi KK, Simons M. The molecular basis of endothelial cell plasticity. Nat Commun. 2017;8:14361. doi: 10.1038/ncomms14361

26. Li Y, Lui KO, Zhou B. Reassessing endothelial-to-mesenchymal transition in cardiovascular diseases. Nat Rev Cardiol. 2018;15:445–456. doi: 10.1038/s41569-018-0023-y

27. Zhan H, Lin CHS, Segal Y, Kaushansky K. The JAK2V617F-bearing vascular niche promotes clonal expansion in myeloproliferative neoplasms. Leukemia. 2018;32:462–469. doi: 10.1038/leu.2017.233

28. Lin CH, Kaushansky K, Zhan H. JAK2(V617F)-mutant vascular niche contributes to JAK2(V617F) clonal expansion in myeloproliferative neoplasms. Blood Cells Mol Dis. 2016;62:42–48. doi: 10.1016/j.bcmd.2016.09.004

29. Yoshihara H, Arai F, Hosokawa K, Hagiwara T, Takubo K, Nakamura Y, Gomei Y, Iwasaki H, Matsuoka S, Miyamoto K, et al. Thrombopoietin/MPL signaling regulates hematopoietic stem cell quiescence and interaction with the osteoblastic niche. Cell Stem Cell. 2007;1:685–697. doi: 10.1016/j.stem.2007.10.020

30. Olson TS, Caselli A, Otsuru S, Hofmann TJ, Williams R, Paolucci P, Dominici M, Horwitz EM. Megakaryocytes promote murine osteoblastic HSC niche expansion and stem cell engraftment after radioablative conditioning. Blood. 2013;121:5238–5249. doi: 10.1182/blood-2012-10-463414

31. Li H, Zhao N, Li Y, Xing H, Chen S, Xu Y, Tang K, Tian Z, Wang M, Rao Q, et al. c-MPL Is a Candidate Surface Marker and Confers Self-Renewal, Quiescence, Chemotherapy Resistance, and Leukemia Initiation Potential in Leukemia Stem Cells. Stem Cells. 2018;36:1685–1696. doi: 10.1002/stem.2897

32. Ghinassi B, Zingariello M, Martelli F, Lorenzini R, Vannucchi AM, Rana RA, Nishikawa M, Migliaccio G, Mascarenhas J, Migliaccio AR. Increased differentiation of dermal mast cells in mice lacking the Mpl gene. Stem Cells Dev. 2009;18:1081–1092. doi: 10.1089/scd.2008.0323

33. Ng AP, Kauppi M, Metcalf D, Hyland CD, Josefsson EC, Lebois M, Zhang JG, Baldwin TM, Di Rago L, Hilton DJ, et al. Mpl expression on megakaryocytes and platelets is dispensable for thrombopoiesis but essential to prevent myeloproliferation. Proc Natl Acad Sci U S A. 2014;111:5884–5889. doi: 10.1073/pnas.1404354111

34. Nishikawa S, Arai S, Masamoto Y, Kagoya Y, Toya T, Watanabe-Okochi N, Kurokawa M. Thrombopoietin/MPL signaling confers growth and survival capacity to CD41-positive cells in a mouse model of Evi1 leukemia. Blood. 2014;124:3587–3596. doi: 10.1182/blood-2013-12-546275

35. Zhan H, Ma Y, Lin CH, Kaushansky K. JAK2V617F-mutant megakaryocytes contribute to hematopoietic stem/progenitor cell expansion in a model of murine myeloproliferation. Leukemia. 2016. doi: 10.1038/leu.2016.114

36. Lin CH, Kaushansky K, Zhan H. JAK2V617F-mutant vascular niche contributes to JAK2V617F clonal expansion in myeloproliferative neoplasms. Blood Cells Mol Dis. 2016;62:42–48. doi: 10.1016/j.bcmd.2016.09.004

37. Zhan H, Ma Y, Lin CH, Kaushansky K. JAK2(V617F)-mutant megakaryocytes contribute to hematopoietic stem/progenitor cell expansion in a model of murine myeloproliferation. Leukemia. 2016;30:2332–2341. doi: 10.1038/leu.2016.114

38. Carpentier G, Berndt S, Ferratge S, Rasband W, Cuendet M, Uzan G, Albanese P. Angiogenesis Analyzer for ImageJ - A comparative morphometric analysis of "Endothelial Tube Formation Assay" and "Fibrin Bead Assay". Sci Rep. 2020;10:11568. doi: 10.1038/s41598-020-67289-8

39. Zacchigna S, Paldino A, Falcao-Pires I, Daskalopoulos EP, Dal Ferro M, Vodret S, Lesizza P, Cannata A, Miranda-Silva D, Lourenco AP, et al. Towards standardization of echocardiography for the evaluation of left ventricular function in adult rodents: a position paper of the ESC Working Group on Myocardial Function. Cardiovasc Res. 2021;117:43–59. doi: 10.1093/cvr/cvaa110

40. Taqueti VR, Di Carli MF. Coronary Microvascular Disease Pathogenic Mechanisms and Therapeutic Options: JACC State-of-the-Art Review. J Am Coll Cardiol. 2018;72:2625–2641. doi: 10.1016/j.jacc.2018.09.042

41. Hiemann NE, Wellnhofer E, Knosalla C, Lehmkuhl HB, Stein J, Hetzer R, Meyer R. Prognostic impact of microvasculopathy on survival after heart transplantation: evidence from 9713 endomyocardial biopsies. Circulation. 2007;116:1274–1282. doi: 10.1161/CIRCULATIONAHA.106.647149

42. Hao Y, Stuart T, Kowalski MH, Choudhary S, Hoffman P, Hartman A, Srivastava A, Molla G, Madad S, Fernandez-Granda C, et al. Dictionary learning for integrative, multimodal and scalable single-cell analysis. Nat Biotechnol. 2024;42:293–304. doi: 10.1038/s41587-023-01767-y

43. Young MD, Behjati S. SoupX removes ambient RNA contamination from droplet-based single-cell RNA sequencing data. Gigascience. 2020;9. doi: 10.1093/gigascience/giaa151

44. McGinnis CS, Murrow LM, Gartner ZJ. DoubletFinder: Doublet Detection in Single-Cell RNA Sequencing Data Using Artificial Nearest Neighbors. Cell Syst. 2019;8:329–337 e324. doi: 10.1016/j.cels.2019.03.003

45. Subramanian A, Tamayo P, Mootha VK, Mukherjee S, Ebert BL, Gillette MA, Paulovich A, Pomeroy SL, Golub TR, Lander ES, et al. Gene set enrichment analysis: a knowledge-based approach for interpreting genome-wide expression profiles. Proc Natl Acad Sci U S A. 2005;102:15545–15550. doi: 10.1073/pnas.0506580102

46. Skelly DA, Squiers GT, McLellan MA, Bolisetty MT, Robson P, Rosenthal NA, Pinto AR. Single-Cell Transcriptional Profiling Reveals Cellular Diversity and Intercommunication in the Mouse Heart. Cell Rep. 2018;22:600–610. doi: 10.1016/j.celrep.2017.12.072

47. Cai H, Harrison DG. Endothelial dysfunction in cardiovascular diseases: the role of oxidant stress. Circ Res. 2000;87:840–844. doi: 10.1161/01.res.87.10.840

48. Kalucka J, de Rooij L, Goveia J, Rohlenova K, Dumas SJ, Meta E, Conchinha NV, Taverna F, Teuwen LA, Veys K, et al. Single-Cell Transcriptome Atlas of Murine Endothelial Cells. Cell. 2020;180:764–779 e720. doi: 10.1016/j.cell.2020.01.015

49. Peisker F, Halder M, Nagai J, Ziegler S, Kaesler N, Hoeft K, Li R, Bindels EMJ, Kuppe C, Moellmann J, et al. Mapping the cardiac vascular niche in heart failure. Nat Commun. 2022;13:3027. doi: 10.1038/s41467-022-30682-0

50. Wu B, Zhang Z, Lui W, Chen X, Wang Y, Chamberlain AA, Moreno-Rodriguez RA, Markwald RR, O’Rourke BP, Sharp DJ, et al. Endocardial cells form the coronary arteries by angiogenesis through myocardial-endocardial VEGF signaling. Cell. 2012;151:1083–1096. doi: 10.1016/j.cell.2012.10.023

51. Tian X, Hu T, Zhang H, He L, Huang X, Liu Q, Yu W, He L, Yang Z, Yan Y, et al. Vessel formation. De novo formation of a distinct coronary vascular population in neonatal heart. Science. 2014;345:90–94. doi: 10.1126/science.1251487

52. Brutsaert DL. Cardiac endothelial-myocardial signaling: its role in cardiac growth, contractile performance, and rhythmicity. Physiol Rev. 2003;83:59–115. doi: 10.1152/physrev.00017.2002

53. Zhang H, Lui KO, Zhou B. Endocardial Cell Plasticity in Cardiac Development, Diseases and Regeneration. Circ Res. 2018;122:774–789. doi: 10.1161/CIRCRESAHA.117.312136

54. Mendez Luque LF, Blackmon AL, Ramanathan G, Fleischman AG. Key Role of Inflammation in Myeloproliferative Neoplasms: Instigator of Disease Initiation, Progression. and Symptoms. Curr Hematol Malig Rep. 2019;14:145–153. doi: 10.1007/s11899-019-00508-w

55. Evans CE, Iruela-Arispe ML, Zhao YY. Mechanisms of Endothelial Regeneration and Vascular Repair and Their Application to Regenerative Medicine. Am J Pathol. 2021;191:52–65. doi: 10.1016/j.ajpath.2020.10.001

56. Bhat FA, Advani J, Khan AA, Mohan S, Pal A, Gowda H, Chakrabarti P, Keshava Prasad TS, Chatterjee A. A network map of thrombopoietin signaling. J Cell Commun Signal. 2018;12:737–743. doi: 10.1007/s12079-018-0480-4

57. Kaser A, Brandacher G, Steurer W, Kaser S, Offner FA, Zoller H, Theurl I, Widder W, Molnar C, Ludwiczek O, et al. Interleukin-6 stimulates thrombopoiesis through thrombopoietin: role in inflammatory thrombocytosis. Blood. 2001;98:2720–2725. doi: 10.1182/blood.v98.9.2720

58. Wang W, Liu W, Fidler T, Wang Y, Tang Y, Woods B, Welch C, Cai B, Silvestre-Roig C, Ai D, et al. Macrophage Inflammation, Erythrophagocytosis, and Accelerated Atherosclerosis in Jak2 (V617F) Mice. Circ Res. 2018;123:e35–e47. doi: 10.1161/CIRCRESAHA.118.313283

59. Sano S, Wang Y, Yura Y, Sano M, Oshima K, Yang Y, Katanasaka Y, Min KD, Matsuura S, Ravid K, et al. JAK2 (V617F) -Mediated Clonal Hematopoiesis Accelerates Pathological Remodeling in Murine Heart Failure. JACC Basic Transl Sci. 2019;4:684–697. doi: 10.1016/j.jacbts.2019.05.013

60. Sofi F, Abbate R, Gensini GF, Casini A. Accruing evidence on benefits of adherence to the Mediterranean diet on health: an updated systematic review and meta-analysis. Am J Clin Nutr. 2010;92:1189–1196. doi: 10.3945/ajcn.2010.29673

61. Collaborators GBDD. Health effects of dietary risks in 195 countries, 1990-2017: a systematic analysis for the Global Burden of Disease Study 2017. Lancet. 2019;393:1958–1972. doi: 10.1016/S0140-6736(19)30041-8

62. Fewkes JJ, Kellow NJ, Cowan SF, Williamson G, Dordevic AL. A single, high-fat meal adversely affects postprandial endothelial function: a systematic review and meta-analysis. Am J Clin Nutr. 2022;116:699–729. doi: 10.1093/ajcn/nqac153

63. Tombor LS, Dimmeler S. Why is endothelial resilience key to maintain cardiac health? Basic Research in Cardiology. 2022;117:35. doi: 10.1007/s00395-022-00941-8

64. Poisson J, Tanguy M, Davy H, Camara F, El Mdawar MB, Kheloufi M, Dagher T, Devue C, Lasselin J, Plessier A, et al. Erythrocyte-derived microvesicles induce arterial spasms in JAK2V617F myeloproliferative neoplasm. J Clin Invest. 2020;130:2630–2643. doi: 10.1172/JCI124566

65. Kimishima Y, Misaka T, Yokokawa T, Wada K, Ueda K, Sugimoto K, Minakawa K, Nakazato K, Ishida T, Oshima M, et al. Clonal hematopoiesis with JAK2V617F promotes pulmonary hypertension with ALK1 upregulation in lung neutrophils. Nat Commun. 2021;12:6177. doi: 10.1038/s41467-021-26435-0

66. Molinaro R, Sellar RS, Vromman A, Sausen G, Folco E, Sukhova GK, McConke ME, Corbo C, Ebert BL, Libby P. The clonal hematopoiesis mutation Jak2(V617F) aggravates endothelial injury and thrombosis in arteries with erosion-like intimas. Int J Cardiol. 2024;409:132184. doi: 10.1016/j.ijcard.2024.132184

67. Pikman Y, Lee BH, Mercher T, McDowell E, Ebert BL, Gozo M, Cuker A, Wernig G, Moore S, Galinsky I, et al. MPLW515L is a novel somatic activating mutation in myelofibrosis with myeloid metaplasia. PLoS Med. 2006;3:e270. doi: 10.1371/journal.pmed.0030270

68. Pardanani AD, Levine RL, Lasho T, Pikman Y, Mesa RA, Wadleigh M, Steensma DP, Elliott MA, Wolanskyj AP, Hogan WJ, et al. MPL515 mutations in myeloproliferative and other myeloid disorders: a study of 1182 patients. Blood. 2006;108:3472–3476. doi: 10.1182/blood-2006-04-018879

69. Beer PA, Campbell PJ, Scott LM, Bench AJ, Erber WN, Bareford D, Wilkins BS, Reilly JT, Hasselbalch HC, Bowman R, et al. MPL mutations in myeloproliferative disorders: analysis of the PT-1 cohort. Blood. 2008;112:141–149. doi: 10.1182/blood-2008-01-131664

70. Sangkhae V, Etheridge SL, Kaushansky K, Hitchcock IS. The thrombopoietin receptor, MPL, is critical for development of a JAK2V617F-induced myeloproliferative neoplasm. Blood. 2014;124:3956–3963. doi: 10.1182/blood-2014-07-587238

71. Marty C, Pecquet C, Nivarthi H, El-Khoury M, Chachoua I, Tulliez M, Villeval JL, Raslova H, Kralovics R, Constantinescu SN, et al. Calreticulin mutants in mice induce an MPL-dependent thrombocytosis with frequent progression to myelofibrosis. Blood. 2016;127:1317–1324. doi: 10.1182/blood-2015-11-679571

72. Araki M, Yang Y, Masubuchi N, Hironaka Y, Takei H, Morishita S, Mizukami Y, Kan S, Shirane S, Edahiro Y, et al. Activation of the thrombopoietin receptor by mutant calreticulin in CALR-mutant myeloproliferative neoplasms. Blood. 2016;127:1307–1316. doi: 10.1182/blood-2015-09-671172

73. Chachoua I, Pecquet C, El-Khoury M, Nivarthi H, Albu RI, Marty C, Gryshkova V, Defour JP, Vertenoeil G, Ngo A, et al. Thrombopoietin receptor activation by myeloproliferative neoplasm associated calreticulin mutants. Blood. 2016;127:1325–1335. doi: 10.1182/blood-2015-11-681932

74. Eisenberg LM, Markwald RR. Molecular regulation of atrioventricular valvuloseptal morphogenesis. Circ Res. 1995;77:1–6. doi: 10.1161/01.res.77.1.1

75. Chen Q, Zhang H, Liu Y, Adams S, Eilken H, Stehling M, Corada M, Dejana E, Zhou B, Adams RH. Endothelial cells are progenitors of cardiac pericytes and vascular smooth muscle cells. Nat Commun. 2016;7:12422. doi: 10.1038/ncomms12422

76. Grego-Bessa J, Luna-Zurita L, del Monte G, Bolos V, Melgar P, Arandilla A, Garratt AN, Zang H, Mukouyama YS, Chen H, et al. Notch signaling is essential for ventricular chamber development. Dev Cell. 2007;12:415–429. doi: 10.1016/j.devcel.2006.12.011

77. Miquerol L, Thireau J, Bideaux P, Sturny R, Richard S, Kelly RG. Endothelial plasticity drives arterial remodeling within the endocardium after myocardial infarction. Circ Res. 2015;116:1765–1771. doi: 10.1161/CIRCRESAHA.116.306476

78. Zhao L, Borikova AL, Ben-Yair R, Guner-Ataman B, MacRae CA, Lee RT, Burns CG, Burns CE. Notch signaling regulates cardiomyocyte proliferation during zebrafish heart regeneration. Proc Natl Acad Sci U S A. 2014;111:1403–1408. doi: 10.1073/pnas.1311705111

79. Zhao L, Ben-Yair R, Burns CE, Burns CG. Endocardial Notch Signaling Promotes Cardiomyocyte Proliferation in the Regenerating Zebrafish Heart through Wnt Pathway Antagonism. Cell Rep. 2019;26:546–554 e545. doi: 10.1016/j.celrep.2018.12.048

80. Kikuchi K, Holdway JE, Major RJ, Blum N, Dahn RD, Begemann G, Poss KD. Retinoic acid production by endocardium and epicardium is an injury response essential for zebrafish heart regeneration. Dev Cell. 2011;20:397–404. doi: 10.1016/j.devcel.2011.01.010

81. Tzoumas A, Tapp D, Crousillat D, Honigberg MC, Rodriguez-Lozano PF, Aggarwal NR, Ebong IA, Briller J, De Oliveira GM, Harrington CM, et al. The Role of Coronary Microvascular Dysfunction in Heart Failure With Preserved Ejection Fraction. JACC Adv. 2025;4:102345. doi: 10.1016/j.jacadv.2025.102345

82. Rehan R, Yong A, Ng M, Weaver J, Puranik R. Coronary microvascular dysfunction: A review of recent progress and clinical implications. Front Cardiovasc Med. 2023;10:1111721. doi: 10.3389/fcvm.2023.1111721

83. Akhiyat N, Lasho T, Ganji M, Toya T, Shi CX, Chen X, Braggio E, Ahmad A, Corban MT, Stewart K, et al. Clonal Hematopoiesis of Indeterminate Potential Is Associated With Coronary Microvascular Dysfunction In Early Nonobstructive Coronary Artery Disease. Arterioscler Thromb Vasc Biol. 2023;43:774–783. doi: 10.1161/ATVBAHA.122.318928

84. Yu B, Roberts MB, Raffield LM, Zekavat SM, Nguyen NQH, Biggs ML, Brown MR, Griffin G, Desai P, Correa A, et al. Supplemental Association of Clonal Hematopoiesis With Incident Heart Failure. J Am Coll Cardiol. 2021;78:42–52. doi: 10.1016/j.jacc.2021.04.085

85. Shi C, Aboumsallem JP, Suthahar N, de Graaf AO, Jansen JH, van Zeventer IA, Bracun V, de Wit S, Screever EM, van den Berg PF, et al. Clonal haematopoiesis of indeterminate potential: associations with heart failure incidence, clinical parameters and biomarkers. Eur J Heart Fail. 2023;25:4–13. doi: 10.1002/ejhf.2715

86. Schuermans A, Honigberg MC, Raffield LM, Yu B, Roberts MB, Kooperberg C, Desai P, Carson AP, Shah AM, Ballantyne CM, et al. Clonal Hematopoiesis and Incident Heart Failure With Preserved Ejection Fraction. JAMA Network Open. 2024;7:e2353244–e2353244. doi: 10.1001/jamanetworkopen.2023.53244

87. Harada T, Sorimachi H, Obokata M, Naser JA, Ibe T, Tada A, Doi S, Kazui S, Murakami T, Kagami K, et al. Echocardiographic Diastolic Function Grading in HFpEF: Testing the Updated 2025 ASE Criteria. J Am Coll Cardiol. 2026. doi: 10.1016/j.jacc.2025.11.024

