## Supplementary figure for "Targeting TPO/MPL Signaling to Mitigate JAK2V617F-driven Cardiac Microvascular Disease"

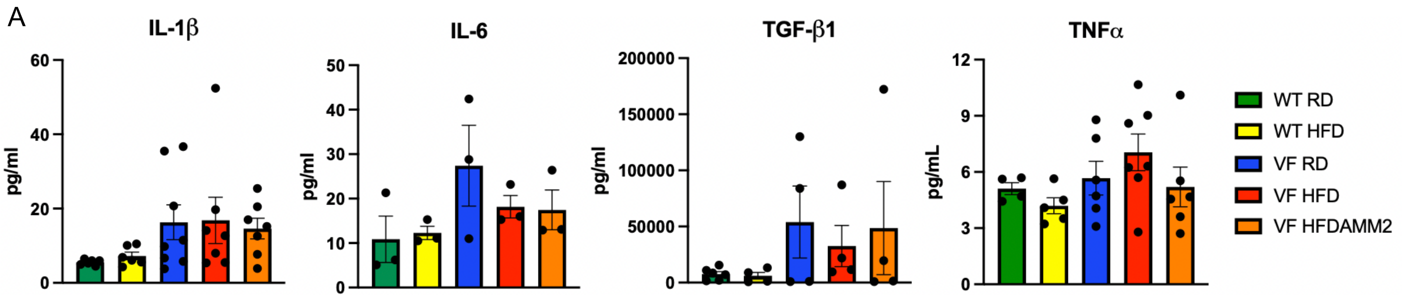


**Supplementary Figure 1.** Plasma levels of IL1β (n=6-8 mice per group), IL6 (n=3 mice per group), TGFβ (n=4-6 mice per group), TNFα (n=4-7 mice per group) proteins in wild-type (WT) and JAK2V617F mutant (VF) mice after 5 weeks of regular diet (RD), HFD, or HFD plus AMM2 treatment.

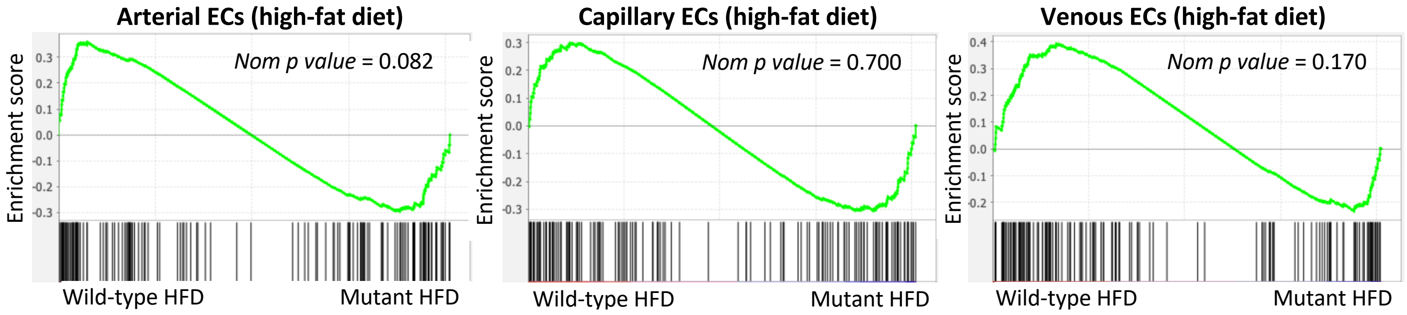


**Supplementary Figure 2.** GSEA of differentially expressed TPO/MPL signaling genes in arterial (left), capillary (middle), and venous (right) ECs in HFD-treated JAK2V617F mutant mice (Mutant HFD) compared with wild-type mice (Wild-type HFD).
